# Discovery of a tRNA-regulatory transcription factor that suppresses breast cancer metastasis

**DOI:** 10.1101/2025.04.26.650725

**Authors:** Siyu Chen, Daniel Markett, Mehran Karimzadeh, Yikai Luo, Matvei Khoroshkin, Baris Boyraz, Sean Lee, Chris Carpenter, Phi Nguyen, Kristle Garcia, Tanvi Joshi, Charlotte Martin, Benjamin Hänisch, Henrik Molina, Sohail Tavazoie, Vijay Ramani, Albertas Navickas, Hani Goodarzi

**Author notes:** Corresponding authors. (HG), (AN), (VR). These Authors contributed equally to this work (Siyu Chen, Daniel Markett).

## Abstract

Transfer RNAs (tRNAs), once viewed as static adaptors in translation, are now recognized as dynamic regulators of gene expression. While recent studies have illuminated roles for tRNA stability, the upstream mechanisms governing tRNA transcription remain poorly understood. To address this gap, we generated the EXpression atlas of tRNA (EXTRNA), a high-resolution tRNA expression dataset spanning 24 cell lines across 9 human tissues. EXTRNA revealed both tissue-type-specific expression programs (“tRNAomes”) and unexpected intra-tissue heterogeneity across breast cancer samples. Integrating EXTRNA with computational network analysis and data from other publicly available datasets, we identified Zinc Finger ZZ-Type And EF-Hand Domain Containing 1 (ZZEF1) as the first sequence-specific transcription factor of a particular tRNA. ZZEF1 promoted tRNA-Lys^UUU^ transcription by partnering with the ATP-dependent chromatin remodeler Chromodomain Helicase DNA Binding Protein 6 (CHD6), enhancing chromatin accessibility at tRNA-Lys-TTT-3 loci. ZZEF1 deficiency reduced tRNA-Lys^UUU^ abundance, decreased the translational efficiency of AAR codon-enriched mRNAs—including the tumor suppressor Serine/Threonine Kinase 3 (STK3)—and promoted metastatic progression in breast cancer *in vivo*. Together, our findings establish a previously unrecognized mechanism for RNA polymerase III–mediated tRNA transcription and define a regulatory circuit linking chromatin remodeling, codon-specific translation, and tumor suppression. More broadly, this work introduces a framework for dissecting the regulatory logic of the tRNAome and highlights tRNA expression control as a promising avenue for therapeutic intervention.

## Introduction

Transfer RNAs (tRNAs) have long been regarded as static adaptor molecules in the central dogma of molecular biology, decoding genetic codons in mRNAs and translating them into amino acids during protein synthesis^1,2^. It is well established that tRNA abundances have coevolved with codon usage across genomes^3^ to optimize protein synthesis^4^. Highly expressed genes frequently use codons decoded by abundant tRNAs, driven by natural selection on translational efficiency and accuracy^5–8^. Perturbation of this balance can destabilize mRNA, impair translational efficiency and accuracy, and subsequently disrupt protein abundance, folding, functional integrity, and overall organismal fitness^9–15^.

Despite their essential role, our understanding of tRNA expression and regulation has suffered due to methodological and technical limitations. In eukaryotes, tRNAs are small (70-90 nucleotides), extensively modified RNAs that adopt conversed secondary and tertiary structures^16–18^. On average, each tRNA molecule harbors 13 covalent modifications, which enhance stability and decoding fidelity during translation^19^. These structural and chemical complexities have historically hindered accurate quantification of tRNAs^20^. For decades, tRNA gene copy number served as a proxy for tRNA abundance^21^. The advent of microarray technologies later enabled systematic measurements, revealing that tRNA expression exhibits tissue-specific patterns^22^. More recently, studies have identified specific tRNA expression profiles associated with disease progression, particularly in shaping gene expression and driving conditions like breast cancer metastasis^20,23,24^. These discoveries challenge the traditional view of tRNAs as passive translation intermediates, and instead position them as critical regulators of disease processes. While these findings establish the functional relevance of specific tRNAs in distinct biological contexts, a systematic and comprehensive understanding of tRNA expression dynamics across different cell types and disease states remains lacking. Generating a large-scale tRNA expression compendium spanning multiple tissues and pathological conditions is essential for uncovering broader regulatory principles and mechanistic insights into tRNA function.

Dynamic tRNA expression patterns across conditions suggest the existence of complex regulatory mechanisms. Several biological processes have been investigated in this context. For instance, tRNA modifications play an important role in maintaining stability and fidelity during translation by influencing proper tRNA folding, accurate aminoacylation by aminoacyl-tRNA synthetases, and precise codon recognition, particularly in wobble base pairing^25,26^. Dysregulated tRNA processing, including the generation of tRNA-derived fragments (tRF), has also been implicated in various diseases^20,27,28^. In cancer cells, oncogenic signaling can alter DNA copy numbers, leading to altered tRNA expression ^24^. Despite these advances, the transcriptional regulation of tRNAs remains largely unexplored.

TRNAs are transcribed by RNA polymerase III (Pol III), a process extensively studied across diverse eukaryotic systems^29–33^. Transcription of tRNA genes begins at internal promoter elements— the A-box and B-box—which are recognized by TFIIIC and subsequently recruit the TFIIIB complex and Pol III^34,35^. While these general mechanisms are well understood, how tRNA transcription is regulated beyond the core Pol III machinery remains unclear. Early *in vitro* studies suggested that upstream flanking regions can influence transcriptional activity^36^, and recent work has shown that nearby RNA polymerase II (Pol II) activity may modulate Pol III transcription^37,38^. The mechanistic target of rapamycin complex 1 (mTORC1) substrate MAF1, a conserved global repressor of Pol III, adds another regulatory layer, but lacks specificity for individual tRNA genes^39–41^. Thus, the functional and sequence-specific determinants of tRNA transcription remain largely undefined—limiting our ability to link transcriptional control to tRNA function in health and disease.

Here, we generated a tRNA-seq dataset from 24 cell lines spanning nine tissue types, revealing tissue-type-specific tRNA expression patterns and substantial intra-tissue heterogeneity. By integrating these data with other publicly available datasets and conducting additional experiments, we identified ZZEF1 as a novel transcriptional regulator of tRNA-Lys^UUU^. Through a combination of computational and experimental approaches, we showed that ZZEF1 binding increases chromatin accessibility at tRNA-Lys-TTT-3 loci, a process facilitated by the chromatin remodeler CHD6. This regulatory axis impacts the translational efficiency of proteins that frequently use AAR codons, including the tumor suppressor STK3, in a codon-dependent manner, ultimately preventing metastatic progression in breast cancer. Furthermore, we identified sequence-specific binding preferences that underpin ZZEF1’s locus-specific regulation. Our study provides a new perspective on the transcriptional regulation of tRNAs, revealing how chromatin remodeling orchestrates tRNA expression and offering critical insights into the broader regulatory landscape of Pol III transcribed non-coding RNAs. Our findings serve as a blueprint for revealing intricate mechanisms underlying tRNA expression, both in the context of normal cell physiology and human disease.

## Results

### TRNA expression exhibits tissue-type-specific pattern and intratumor heterogeneity

Although tRNAs are increasingly recognized as regulatory molecules^20^, a systematic understanding of their expression variation in disease-relevant physiological contexts across human tissues is still lacking. To fill this gap, we generated an EXpression atlas of tRNA (EXTRNA), a tRNA-seq dataset from 24 cell lines spanning 9 tissue types (**Fig. 1A**), representing diverse cancer types from the Cancer Cell Line Encyclopedia (CCLE)^42^. To systematically characterize tRNA expression landscapes across these cell lines, we performed a series of atlas-wide comparisons.

**Figure 1.**
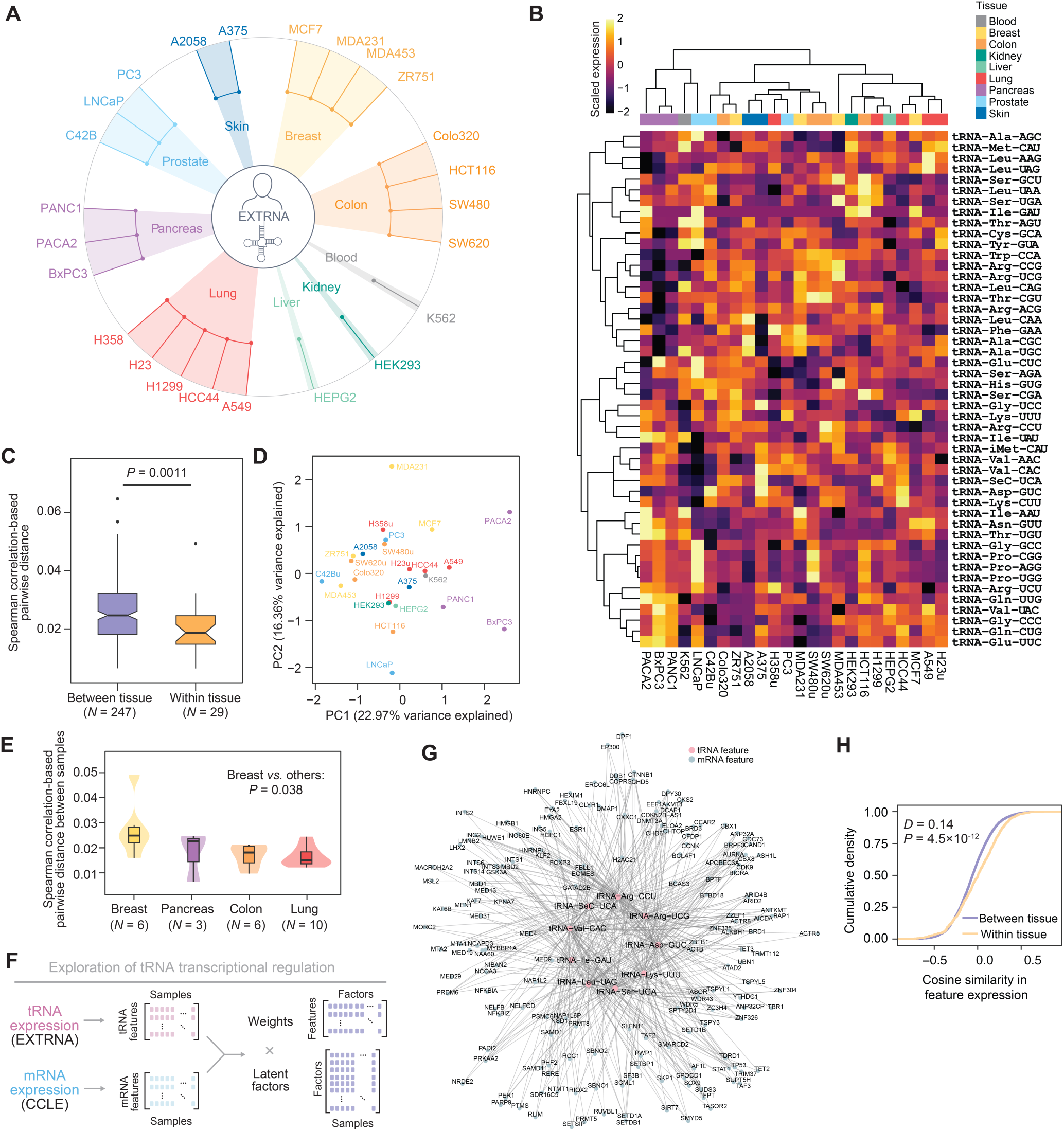
Dynamic and heterogeneous expression of tRNAs revealed by the EXTRNA dataset. **(A)** Schematic illustration of EXTRNA, a large compendium of tRNA expression profiles across 24 cell lines representing 9 tissue types. **(B)** Heatmap displaying tRNA isoacceptor expression profiles obtained in EXTRNA. Hierarchical clustering was applied to group the profiles. **(C)** Boxplot comparing Spearman correlation-based sample-sample pairwise distances calculated from tRNA isoacceptor expression profiles between tissue types and within tissue types. *P*-value was calculated using the Mann– Whitney *U* test. **(D)** Principal Component Analysis (PCA) plot visualizing tissue-type-specific clustering patterns in tRNA expression profiles. **(E)** Violin and boxplot representation of Spearman correlation-based sample-sample pairwise distances calculated from tRNA isoacceptor expression profiles within tissue types. Tissues are ordered by the median distance values. *P*-value was calculated using the Kolmogorov–Smirnov test. **(F)** Schematic representation of the joint analysis integrating tRNA isoacceptor expression from EXTRNA with RNA expression data from CCLE to explore regulatory factors of tRNA transcription. **(G)** Example of a constructed transcriptional network from MOFA+ joint analysis, illustrating regulatory interactions identified within a representative module (factor 4, cluster 6). **(H)** Cumulative density curves showing the expression similarity of tRNA and RNA features among cell lines within the same tissue type and between different tissue types. *D*-value and *P*-value were calculated using the Kolmogorov–Smirnov test.

Expression of both tRNA isoacceptors (which decode synonymous codons for the same amino acid) and isodecoders (which share anticodons but differ in body sequence) varied substantially across cell lines, revealing strong cell-type specificity (**Fig. 1B**, **S1A** and **S1B**). This observation is consistent with our previous work showing that tRNA isoacceptors have functionally diverging roles^23^. Beyond overall variability, tRNA expression profiles were more similar among cell lines from the same tissue type than across different tissues (**Fig. 1C**, *P* = 0.001, Mann–Whitney *U* test). Principal component analysis (PCA) further supported this trend, showing significant tissue-type clustering (**Fig. 1D** and **S1C**, *P* = 0.04, MANOVA test). These results suggest that tRNAs exhibit tissue-type-specific expression patterns.

In addition to global tissue-type specificity, specific tissues exhibited notable heterogeneity in PCA space (**Fig. S1D**, *P* = 0.046, Kruskal–Wallis test). To quantify this intra-tissue variation, we computed pairwise Spearman-correlation-based distances between genetically distinct cell lines within each tissue sampled by EXTRNA, providing a direct measure of intra-tissue variability in tRNA isoacceptor expression (**Fig. 1E**). Notably, breast cancer cell lines displayed particularly high heterogeneity (**Fig. 1E** and **S1E**, *P* = 0.038, Kolmogorov–Smirnov test), consistent with our previous findings that tRNA isoacceptor expression varies in breast cancer^23,24^. Together, this atlas and these results imply the existence of tissue-, tissue-subtype, and cancer-specific regulatory programs for tRNA expression.

### Context-dependent transcriptional regulation underlies tRNA expression variation

The regulatory basis of tRNA expression variation across tissues and cancer types remains poorly understood, particularly at the level of transcriptional control^20^. To address this, we investigated transcription-associated factors potentially linked to the context-specific expression patterns observed in EXTRNA. Specifically, we employed MOFA+, a multi-modal data integration framework^43^, to integrate tRNA isoacceptor expression from EXTRNA with mRNA expression data from the CCLE dataset (**Fig. 1F**). We focused our covariation analysis on genes involved in transcriptional regulation, chromatin remodeling, and histone binding. The variance explained by each latent factor (**Fig. S1F**) and the distribution of factor values across samples (**Fig. S1G**) obtained from MOFA+ analysis indicate complex associations between the expression of transcriptional regulators and specific tRNA species. Latent factor values were generally more similar in a given tissue than across tissues (**Fig. S1G** and **S1H**, *P* = 0.034, Mann–Whitney *U* test), indicating that tissue-dependent transcriptional regulation underlie the observed tissue-type-specific tRNA expression patterns (e.g. factors 2 and 5 significantly associating with tissue type; **Fig. S1I**; *P* = 0.04 and 0.01 for factor 2 and 5, respectively, ANOVA test). However, we note several other latent factors capturing variation beyond tissue identity (**Fig. S1I**), implying that tRNA expression is further shaped by regulatory programs outside tissue-specific contexts.

To further dissect the regulatory architecture underlying MOFA+ associations, we applied *k*-means clustering to identify interactions between co-expressed transcriptional regulators and tRNAs within each latent factor. Clustering analyses revealed structured regulatory modules—groups of tRNAs and regulators that strongly co-vary across the EXTRNA dataset (**Fig. 1G** and **S1J**).

Reassuringly, these modules contained well-established transcriptional regulators of Pol II activity, including FOXP3, JUN, and CDK9; these hits align with prior studies linking Pol II activity to Pol III regulation^37,38^. In addition, we observed multiple chromatin-associated regulators covarying with the expression of specific tRNAs, including ATP-dependent chromatin remodeling enzymes (e.g. CHD6), lysine deacetylases (e.g. SIRT2), and putative transcription regulators (e.g. ZZEF1). These modules nominate potential chromatin-associated regulators that may contribute to the context-specific tRNA expression.

Given our observation of both tissue-type specificity and intra-tissue heterogeneity in tRNA expression, we next sought to evaluate whether the structured regulatory modules reflect these patterns and help explain the underlying transcriptional regulation. Specifically, we quantified (1) tissue specificity and (2) intra-tissue heterogeneity of each module by calculating the pairwise cosine similarity of tRNA and mRNA expression profiles across cell lines (e.g., **Fig. 1G** and **S1J**). Consistent with the tissue-specific patterns observed in tRNA expression, cell lines from the same tissue type exhibited significantly higher expression similarity within modules than those from different tissues (**Fig. 1H**, *P* = 4.5×10^-12^, Kolmogorov–Smirnov test). To assess heterogeneity within tissues, we calculated the variance in expression similarity among cell lines within each tissue. Notably, breast cancer cell lines exhibited significantly higher intra-tissue variability than other tissues (**Fig. S1K**, breast *vs.* lung, *P* = 0.006; breast *vs.* pancreas, *P* = 2×10^-4^, Mann–Whitney *U* test). Together, these results indicate that tRNA transcriptional regulation is more complex than previously appreciated: it is context-dependent, and shaped by a mixture of tissue-intrinsic and broadly acting chromatin-associated regulators.

### ZZEF1 functions as a regulator of tRNA-Lys^UUU^ transcription in breast cancer

Our pan-cancer analysis revealed highly context-dependent regulatory programs underlying tRNA expression, with breast cancer cell lines displaying particularly high intra-tissue heterogeneity. To further dissect transcriptional regulation in this context, we focused on identifying breast cancer– specific master regulators of tRNA expression. Master regulator analysis requires a large number of independent samples to reliably infer regulatory interactions, EXTRNA alone, while providing valuable insights into tRNA expression variation, is not sufficient on its own to support such analysis. To overcome this limitation, we leveraged The Cancer Genome Atlas Breast Cancer (TCGA-BRCA) dataset^44,45^, one of the largest and most comprehensive publicly available collections of breast cancer gene expression measurements. Specifically, we relied on small-RNA sequencing data for these samples, which can be used to infer tRNA expression using tRNA fragment (tRF) quantification as a proxy^44–46^, enabling large-scale analysis of covarying tRFs and their candidate regulators in breast cancer.

To identify master regulators with TCGA-BRCA data, we developed a multi-modal integrative framework incorporating mRNA and tRF expression data^47^ (**Fig. 2A**). In particular, we constructed a unified transcriptional network linking protein-coding genes (measured by mRNA-seq) and tRFs based on their co-expression dependencies across breast cancer samples, using the ARACNe algorithm^48,49^, followed by application of the data processing inequality to eliminate indirect interactions^49^. From the refined network, we identified candidate transcriptional regulators via master regulator analysis^50^, prioritizing factors with nucleic acid-binding capacity and/or nuclease activity (**Fig. 2A** and **S2A**). To validate these candidates, we used CRISPR interference (CRISPRi) to knock down 24 high-confidence regulators in the triple-negative breast cancer cell line MDA-MB-231 (hereafter, “MDA”). Of these, 11 achieved knockdown efficiencies ≥80% and were selected for tRF abundance profiling by small RNA sequencing (**Fig. S2B**). Consistent with our network analysis, we observed significant changes in the abundance of specific tRFs following candidate knockdowns (**Fig. 2B**); knockdown of ZZEF1 altered the abundance of tRFs derived from tRNA-Arg^ACG^, tRNA-Arg^UCG^, tRNA-Thr^AGU^, and tRNA-Lys^UUU^ (**Fig. 2C**); knockdown of ZFP36 significantly altered the abundance of tRFs deriving from tRNA-Asp^GUC^, tRNA-Asn^GUU^, tRNA-Leu^AAG^, tRNA-Thr^AGU^ (**Fig. S2C**); knockdown of ZNF703 knockdown altered the levels of tRFs for tRNA-iMet^CAU^, tRNA-Ser^AGA^, tRNA-Glu^CUC^ and tRNA-Leu^AAG^ (**Fig. S2D**). These results suggest specific regulatory pathways for tRF abundance, and possibly tRNAs themselves.

**Figure 2.**
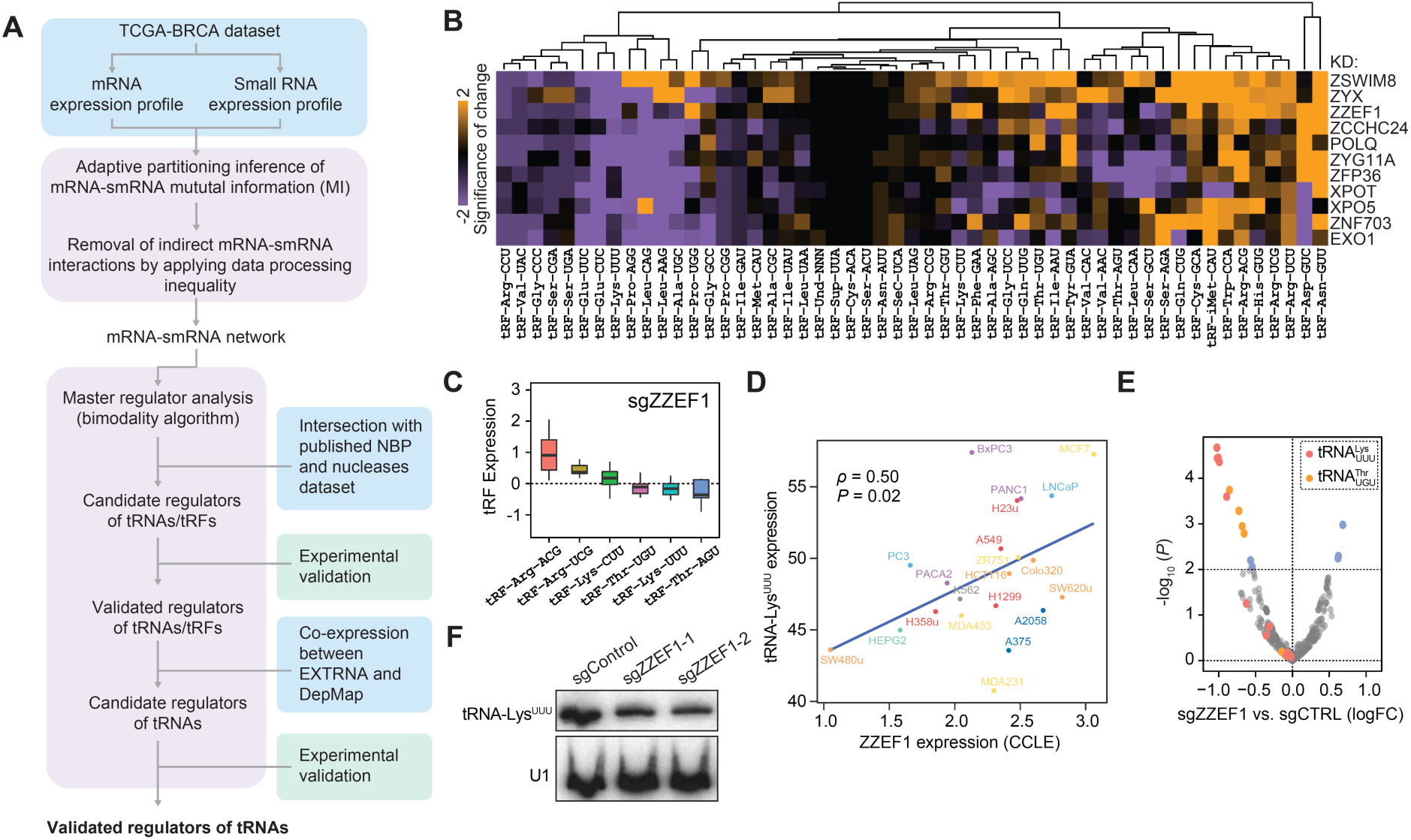
Identification of ZZEF1 as a key regulator of tRNA-Lys^UUU^ in breast cancer cells. **(A)** Schematic representation of the workflow integrating network analysis and experimental validation to identify transcriptional regulators of tRNA expression in breast cancer cells. NBP, nucleic acid-binding proteins. **(B)** Heatmap displaying tRF expression profiles in MDA cells after knockdown of identified transcriptional regulator candidates. Hierarchical clustering was applied to group the resulting profiles. **(C)** tRF expression changes in MDA cells following ZZEF1 knockdown, compared to control cells. **(D)** Correlation analysis showing a significant positive association between tRNA-Lys^UUU^ abundance and ZZEF1 expression across different cancer cell lines. tRNA-Lys^UUU^ abundance was calculated as the sum of the corresponding isodecoders. Spearman’s *ρ* and the associated *P*-value are shown. **(E)** Mature tRNA expression alterations in MDA cells with ZZEF1 knockdown, highlighting significant changes in tRNA-Lys^UUU^ compared to control cells. tRNA molecules with significant abundance alterations (*P* < 0.01) after ZZEF1 knockdown are colored in blue. tRNA-Lys^UUU^ isodecoders are colored in orange, and tRNA-Thr^UGU^ is colored in yellow. **(F)** Northern blot analysis of tRNA-Lys^UUU^ and U1 RNA (loading control) in MDA cells with ZZEF1 knockdown and control cells.

However, the observed impact on tRF abundance alteration can be confounded by tRNA processing and stability. To address this, we leveraged the EXTRNA dataset to evaluate candidate regulators using direct measurements of mature tRNA expression, as we expect that a direct regulator would exhibit co-expression with mature tRNAs across cell lines. Intersecting hits with our EXTRNA and TCGA analyses (**Fig. S2E**) converged on a notable hit with limited prior functional characterization: ZZEF1. ZZEF1 and tRNA-Lys^UUU^ levels positively correlated across cancer cells (**Fig. 2D** and **S2F**, *ρ* = 0.50, *P* = 0.02, Spearman correlation), suggesting that ZZEF1 may serve as a direct positive regulator of tRNA transcription.

To confirm whether ZZEF1 is a direct regulator of tRNA-Lys^UUU^ and the specificity of ZZEF1 for tRNA expression, we performed mature tRNA-seq following ZZEF1 knockdown in MDA cells, including other high-scoring and low-scoring regulatory candidates validated in the previous step (**Fig. 2B** and **S2**). Consistent with our predictions, ZZEF1 knockdown resulted in significant alterations in mature tRNA expression compared to control cells, with notable effects on tRNA-Lys^UUU^ (**Fig. 2E** and **S2G**). We also confirmed this finding using Northern blot analysis (**Fig. 2F**). Together, these results establish ZZEF1 as a critical regulator of tRNA-Lys^UUU^ expression, providing further support for its role in shaping tRNA dynamics in breast cancer.

### ZZEF1-tRNA-Lys^UUU^ axis modulates translation in a codon-dependent manner

An altered tRNA pool can affect protein production through translation^10^. To investigate whether ZZEF1-mediated regulation of tRNA-Lys^UUU^ impacts gene expression at the translational level, we analyzed translational landscapes at both the gene and codon levels using ribosome profiling (Ribo-seq) in ZZEF1 knockdown and control cells, as well as in cells complemented by ectopic overexpression of tRNA-Lys^UUU^ or a control vector. The high quality of our Ribo-seq data was indicated by strong three-nucleotide periodicity in coding regions and fragment lengths predominantly centered at 33–34 nucleotides, supporting the reliability of our data for precise investigation of translation dynamics (**Fig. S3A-C**).

We observed that a substantial number of genes showed altered translational efficiency following ZZEF1 knockdown (**Fig. 3A**), suggesting that ZZEF1 dysregulation disrupts the translational landscape and affects protein output. Notably, proteins that frequently use AAR codons (AAA and AAG, encoding lysine and decoded by tRNA-Lys^UUU^) exhibited significantly reduced translational efficiency after ZZEF1 knockdown (**Fig. 3B** and **S3D**). This result suggests that ZZEF1 modulates translation in a codon-dependent manner mediated by tRNA-Lys^UUU^. Consistently, increasing tRNA-Lys^UUU^ supply in the ZZEF1-deficient background restored the translational efficiency of AAR codon-rich proteins (**Fig. 3C**). To further quantify the impact of the ZZEF1-tRNA-Lys^UUU^ axis on translation, we combined data on translational efficiency changes in ZZEF1 knockdown cells with those from tRNA-Lys^UUU^ overexpression in the same background. We found that proteins with high AAR codon usage (top 20%) exhibited significantly greater changes in translational efficiency compared to other proteins (**Fig. 3D**, *P* = 6×10^-15^, Mann–Whitney *U* test).

**Figure 3.**
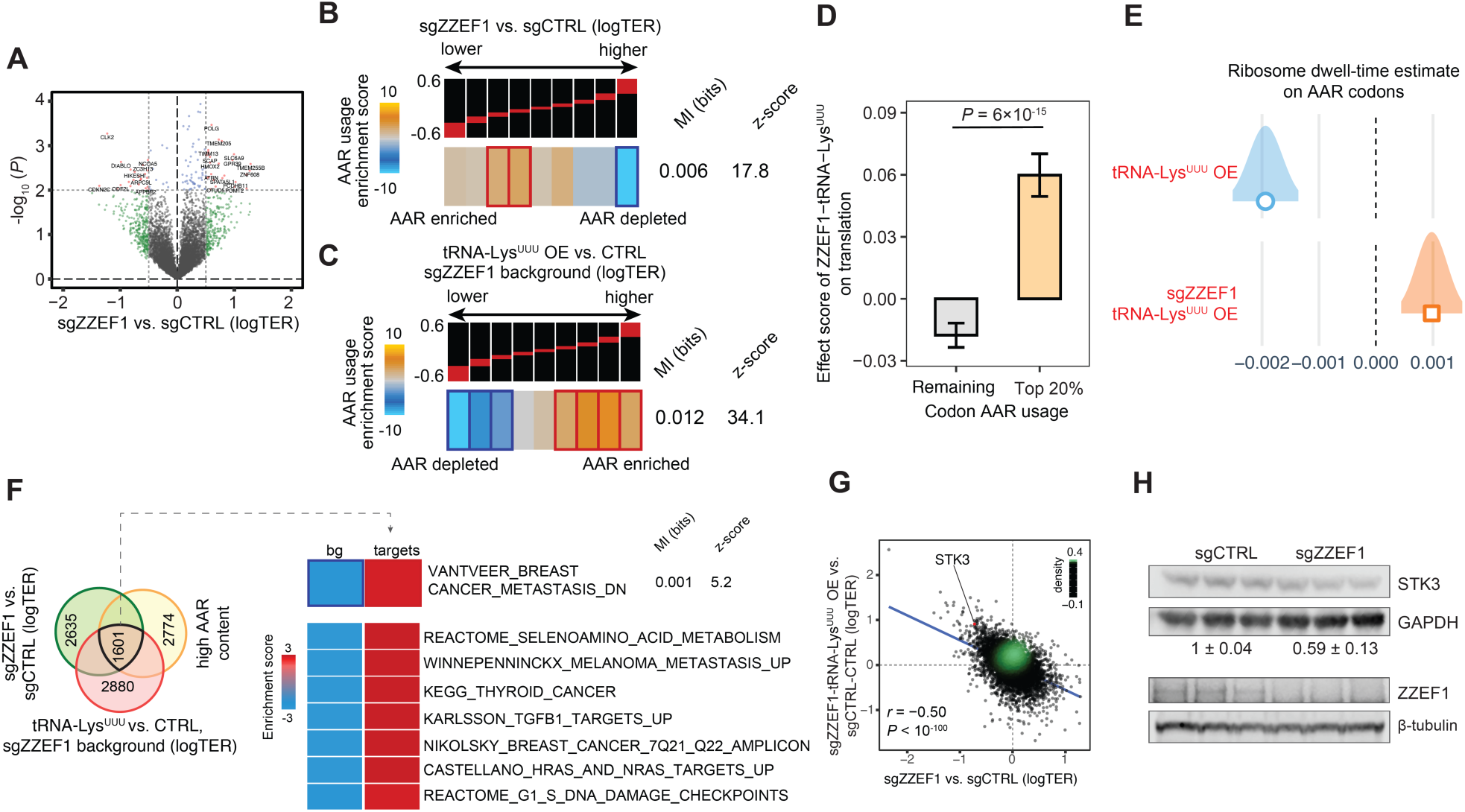
Translation regulation by ZZEF1-tRNA-Lys^UUU^ through codon-dependent mechanisms. **(A)** Volcano plot showing alterations in translational efficiency in MDA cells following ZZEF1 knockdown compared to control cells. TER, translational efficiency ratio. Proteins with an absolute logTER greater than 0.5 and *P* > 0.01 are labeled in green; proteins with an absolute logTER less than 0.5 and *P* < 0.01 are labeled in blue; and proteins with significant logTER alterations (absolute logTER > 0.5 and *P* < 0.01) are colored in red. **(B)** Enrichment of the lysine (AAR codons) content as a function of logTER between ZZEF1 knockdown and control cells. mRNAs are distributed into equally populated bins according to their logTER (the red bars on the black background show the range of values in each bin). Bins with significant enrichment (hypergeometric test, corrected *P* < 0.05; red) or depletion (blue) of AAR codons are denoted with a bolded border. Also included are mutual information (MI) value and its associated z-score. **(C)** Similar to **(B)**, but showing results of enrichment patterns of transcripts with high AAR codon content after restoration of tRNA-Lys^UUU^ supply. **(D)** Bar plot showing transcripts with higher AAR codon content exhibited elevated ZZEF1-tRNA-Lys^UUU^ translational effect scores. **(E)** Ribosomal AAR codon dwelling times estimated using Consistent Excess of Loess Predictions (CELP) bias coefficients in ZZEF1-deficient cells with tRNA-Lys^UUU^ overexpression compared to controls. Higher coefficients indicate longer dwelling times. **(F)** Functional characterization of ZZEF1-tRNA-Lys^UUU^ targets obtained by overlapping proteins that (i) showed significant translational downregulation following ZZEF1 knockdown, (ii) showed significant translational upregulation after tRNA-Lys^UUU^ overexpression in ZZEF1-deficient cells, and (iii) frequently utilized AAR codons. **(G)** Scatter plot showing that proteins with reduced translational efficiency after ZZEF1 knockdown exhibited higher translational efficiency following tRNA-Lys^UUU^ overexpression. Pearson’s *r* and the associated *P*-value are shown. **(H)** Western blot analysis of GAPDH (control) and STK3 in MDA cells with ZZEF1 knockdown compared to control cells (*P* = 0.038). Mean values and standard errors of STK3 quantification are shown. *P*-value was calculated with one-tailed *t*-test.

Together, these results strongly support that ZZEF1-tRNA-Lys^UUU^ modulates translation in a codon-dependent manner.

To further ascertain whether this regulation operates directly at the codon level, we hypothesized that ZZEF1 deficiency would lead to increased ribosome dwelling time on AAR codons due to a reduced translational elongation rate. Using our Ribolog suite^51^, we quantified ribosome dwelling times on AAR codons in ZZEF1-deficient cells with tRNA-Lys^UUU^ overexpression. As predicted, ribosome dwelling times on AAR codons were significantly increased in ZZEF1 knockdown cells (**Fig. 3E**). These results provide direct evidence that the ZZEF1-tRNA-Lys^UUU^ axis regulates translation by modulating ribosome dynamics in a codon-dependent manner.

### Deficiencies of ZZEF1-tRNA-Lys^UUU^-STK3 promote breast cancer metastasis *in vivo*

TRNAs are increasingly recognized as critical regulators in disease development, including cancer progression^10,20^. To investigate the potential phenotypic effects of ZZEF1-tRNA-Lys^UUU^ dysregulation, we first analyzed the functional characteristics of target proteins modulated by the ZZEF1-tRNA-Lys^UUU^ regulatory axis. Specifically, we identified proteins that (i) exhibited significant translational downregulation following ZZEF1 knockdown, (ii) showed significant translational upregulation after increased tRNA-Lys^UUU^ supply in ZZEF1-depleted cells, and (iii) frequently utilized AAR codons. Strikingly, we found that these regulatory targets were enriched in factors associated with tumorigenic and metastatic processes, particularly proteins implicated in breast cancer metastasis (**Fig. 3F**). Among the target proteins, we identified STK3, a well-known tumor suppressor, as one of the top AAR codon users, indicating its regulation as a downstream target of the ZZEF1-tRNA-Lys^UUU^ axis (**Fig. 3G**). This downstream relationship was further validated by western blot analysis (**Fig. 3H**, *P*-value = 0.038, one-tail *t*-test).

To directly evaluate the role of ZZEF1-tRNA-Lys^UUU^-STK3 dysregulation in breast cancer development, we employed xenograft lung colonization assays. In particular, we injected MDA cells with tRNA-Lys^UUU^ knockdown or control intravenously into NOD *scid* gamma (NSG) mice. *In vivo* bioluminescence imaging revealed that tRNA-Lys^UUU^ deficiency significantly increased lung colonization capacity (**Fig. 4A**, *P* = 0.03, ANOVA test), suggesting that tRNA-Lys^UUU^ deficiency promotes breast cancer metastasis. Remarkably, overexpression of tRNA-Lys^UUU^ in MDA cells reduced metastatic colonization compared to controls (**Fig. 4B**, *P* = 0.02, ANOVA test), highlighting the central role of tRNA-Lys^UUU^ in metastasis regulation. To further confirm the contribution of the ZZEF1-tRNA-Lys^UUU^-STK3 deficiency to drive metastasis, we assessed metastatic capacity in xenograft models of control, ZZEF1 knockdown and STK3 knockdown cells using both bioluminescence imaging (**Fig. 4C**) and gross histology of extracted lungs (**Fig. S4A**). As expected, ZZEF1 and STK3 knockdown significantly increased metastatic burdens in the lung (**Fig. 4C** and **S4A**; **Fig. 4C**, *P* = 0.05 and 1×10^-4^ for ZZEF1 knockdown and STK3 knockdown, respectively, ANOVA tests). These results underscore the critical role of the ZZEF1-tRNA-Lys^UUU^-STK3 axis in driving breast cancer metastasis. Additionally, we reproduced these findings in another triple-negative breast cancer cell line, HCC1806 (**Fig. S4B**, *P* < 1×10^-5^ and 0.05 for ZZEF1 knockdown and STK3 knockdown, respectively, ANOVA tests), further indicating the broad relevance of this regulatory axis in breast cancer metastasis.

**Figure 4.**
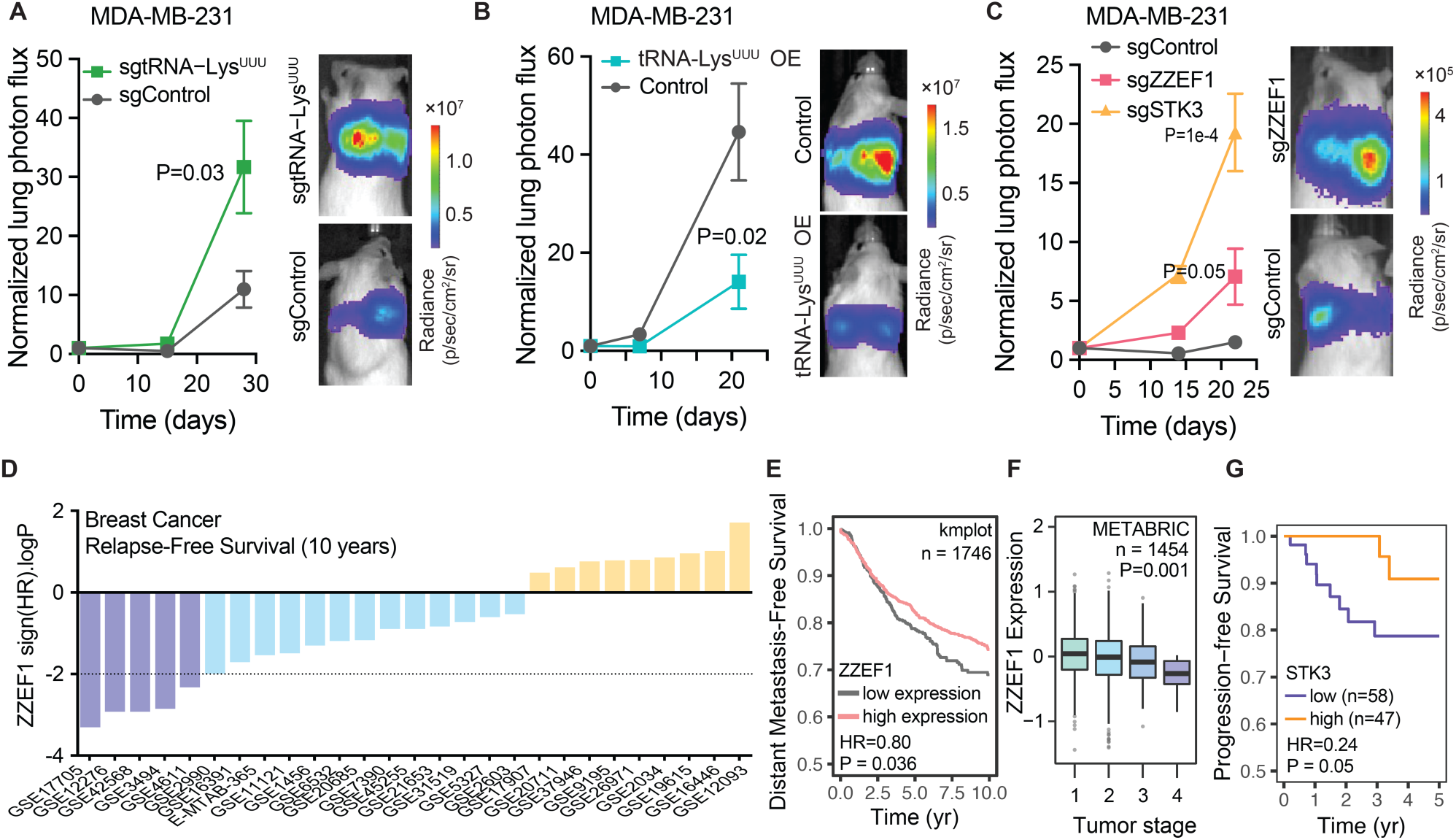
ZZEF1-tRNA-Lys^UUU^-STK3 deficiency promotes metastatic breast cancer and is associated with poor clinical outcomes. **(A)** Bioluminescence imaging showing increased lung colonization in MDA cells with tRNA-Lys^UUU^ knockdown compared to control cells. *N* = 5 mice per cohort. *P* value was calculated using a two-way analysis of variance (ANOVA). **(B)** Similar to **(A)**, but showing reduced lung colonization in MDA cells with tRNA-Lys^UUU^ overexpression compared to control cells. *N* = 5 mice per cohort. **(C)** Bioluminescence imaging showing increased lung colonization in MDA cells with ZZEF1 knockdown and STK3 knockdown compared to control cells. *N* = 4 mice per cohort. **(D)** Bar plot showing the distribution of 10-year relapse-free survival *P*-values (two-sided log-rank test results, reported as –log*P* for positive association and log*P* for negative association) for the correlation between ZZEF1 expression and clinical outcomes across listed breast cancer datasets. Purple bars show associations that pass the statistical threshold (*P* < 0.01). **(E)** Kaplan-Meier survival curve illustrating the correlation between tumor ZZEF1 levels and distant metastasis-free survival in a collection of breast cancer patient cohorts. **(F)** Boxplot showing ZZEF1 expression levels across clinical samples, including tissues from various breast tumor stages. ANOVA was used to calculate the *P*-value. **(G)** Kaplan-Meier survival curve illustrating the correlation between tumor STK3 protein levels and progression-free survival.

To further explore the routes by which the ZZEF1-tRNA-Lys^UUU^-STK3 regulatory axis influences breast cancer metastasis, we conducted proliferation and migration assays using control and ZZEF1 or STK3-deficient MDA cells. Consistent with the *in vivo* results (**Fig. 4C** and **S4A**), knockdown of ZZEF1, or STK3 significantly increased migration (**Fig. S4C,** *P* = 0.005 and 0.037 for ZZEF1 knockdown and STK3 knockdown, respectively, *t*-test) but led to reduced proliferation rates (**Fig. S4D**, *P* = 1×10^-3^ and 3×10^-4^ for ZZEF1 knockdown and STK3 knockdown, respectively, *t*-test). These findings suggest that the ZZEF1-tRNA-Lys^UUU^-STK3 axis suppresses breast cancer metastasis primarily by enhancing cell migration.

### ZZEF1-tRNA-Lys^UUU^-STK3 deficiency is associated with clinical outcomes in breast cancer patients

To assess the clinical relevance of the ZZEF1-tRNA-Lys^UUU^-STK3 axis in breast cancer metastasis, we analyzed publicly available datasets from breast cancer patients. In both individual cohorts and meta-analyses, lower ZZEF1 expression in breast tumors was significantly associated with reduced overall survival, disease-free survival, and distant metastasis-free survival (**Fig. 4D-E** and **S4E-H**). These findings suggest that ZZEF1 expression plays a protective role against tumor progression and metastasis.

Our further analysis revealed an inverse correlation between ZZEF1 expression and tumor stage in breast cancer (**Fig. 4F**), supporting its function as a metastasis suppressor. When stratified by subtype, lower ZZEF1 expression was significantly associated with poorer disease-free survival in HER2+ and basal-like breast tumors (**Fig. S4I**). Additionally, in line with STK3 being a key downstream target of the ZZEF1-tRNA-Lys^UUU^ axis, we found that lower STK3 protein levels were significantly associated with reduced progression-free survival in breast cancer patients (**Fig. 4G**).

Together, these results establish the clinical relevance of the ZZEF1-tRNA-Lys^UUU^-STK3 regulatory axis in suppressing breast cancer metastasis and highlight its potential as a therapeutic target.

### ZZEF1 recognizes distinct sequence features proximal to tRNA loci

Transcriptional regulators typically interact with other factors to exert gene-specific regulatory effects^52^. However, factors that specifically regulate tRNA transcription remain poorly defined. We next thought to investigate the mechanism by which ZZEF1 regulates tRNA-Lys^UUU^ expression. We hypothesized that ZZEF1 directly binds tDNAs encoding tRNA-Lys^UUU^ (i.e. tRNA-Lys-TTT loci). To test it, we performed chromatin immunoprecipitation followed by sequencing (ChIP-seq) in MDA cells using a ZZEF1-specific antibody and an IgG control antibody. We observed significantly enriched ZZEF1 binding compared to control IgG in the vicinity of the tRNA-Lys-TTT-3 loci (hereafter, “target loci”), with substantially higher levels at target loci compared to non-target tRNA-Lys-TTT isodecoder loci, or other tRNA-Lys isoacceptor loci (**Fig. 5A**, **5B** and **S5**). This suggests that ZZEF1 directly binds target tRNA-Lys-TTT-3 isodecoder loci.

**Figure 5.**
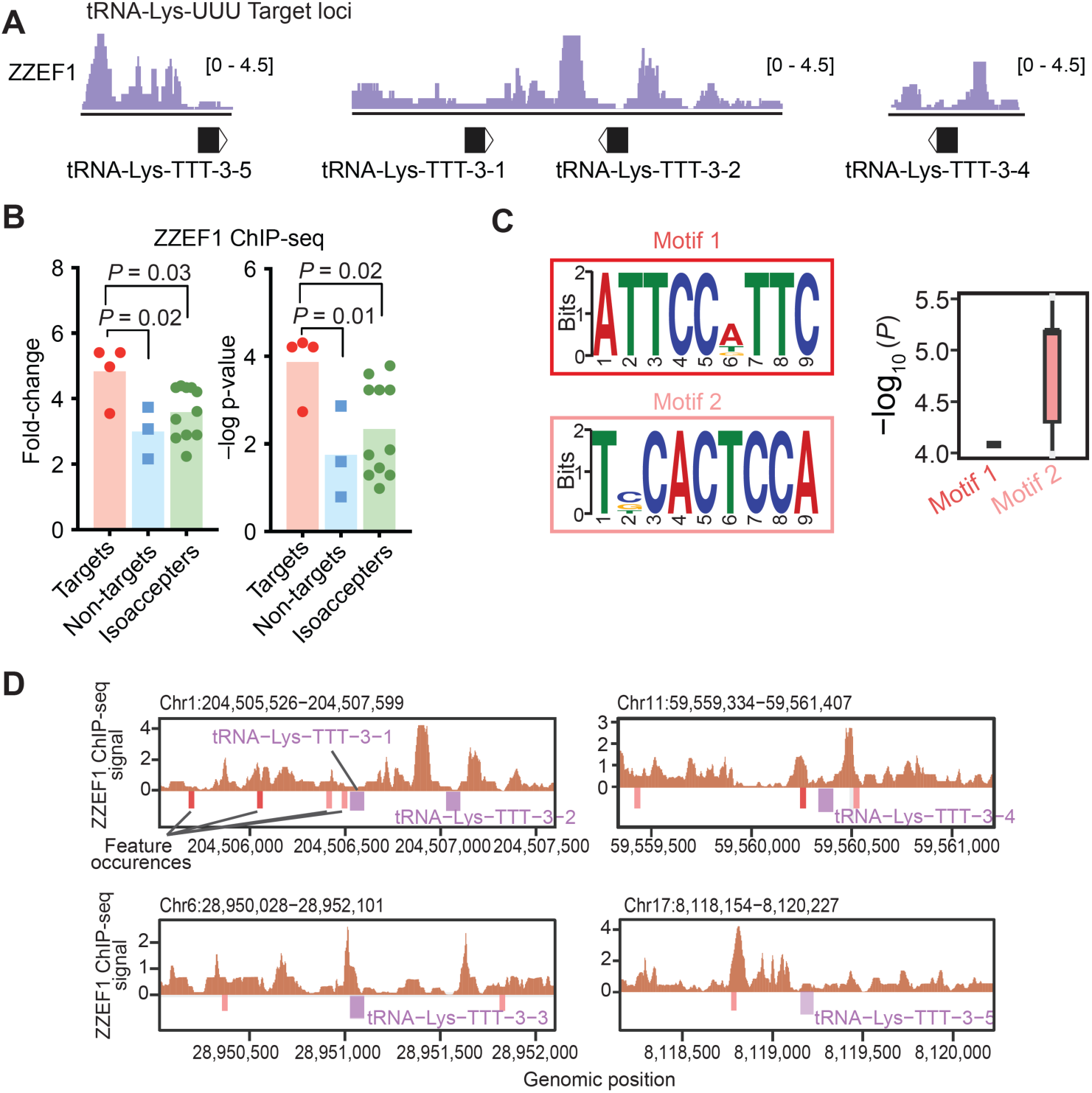
ZZEF1 specifically binds in the proximal regions of tRNA-Lys-TTT-3 loci. **(A)** Normalized ChIP-seq tracks showing enriched ZZEF1 binding in the proximal regions of tRNA-Lys^UUU^-3 loci using a ZZEF1-specific antibody compared to an IgG control in MDA cells. **(B**) Bar plot presenting statistical analysis of ZZEF1 binding enrichment across target loci, non-target loci, and other isoacceptor loci. *P* value calculated using one-tailed Welch’s *t*-test. **(C)** Sequence logo depicting two distinct sequence features that define ZZEF1 binding preferences. Bar plot showing the statistical enrichment (*P*-value) of the identified sequence features in ZZEF1-specific ChIP-seq data compared to genome-wide sequences. **(D)** Schematic representation of the identified sequence features located in the proximal regions of tRNA-Lys-TTT-3 loci.

To understand how ZZEF1 achieves this binding specificity, we hypothesized that, like many transcription factors, ZZEF1 may recognize particular DNA motifs near its targets. To explore this possibility, we performed *de novo* motif discovery^53^ on ZZEF1-enriched binding profiles from ChIP-seq data, and identified two significantly over-represented motifs in ZZEF1 ChIP-seq data (**Fig. 5C**). Notably, these motifs were specifically located in the proximal regions of tRNA-Lys-TTT-3 loci (**Fig. 5D**). These findings suggest that ZZEF1 mediates locus-specific regulation of tRNA-Lys^UUU^ transcription by recognizing distinct sequence elements.

### ZZEF1 promotes tRNA-Lys^UUU^ transcription by increasing chromatin accessibility

Certain transcription factors can bind directly to nucleosomes and act as pioneer factors^54^. Given that ZZEF1 has been characterized as a histone reader^55^, we hypothesized that ZZEF1 might interact directly with nucleosomes to regulate tRNA-Lys^UUU^ expression. To test it, we first examined the interaction between ZZEF1’s domains and nucleosomes *in vitro*. Specifically, we purified poly histidine tag and maltose binding protein (His-MBP) fusion proteins from *Escherichia coli*, including individual domains (EF hand, DOC, ZZ1, ZZ2, and the combined ZZ1/2) as well as MBP alone as a control (**Fig. S6A-B**). We then conducted nucleosome pulldown assays using recombinant mononucleosomes with biotinylated DNA, followed by western blotting for the ZZEF1 domain fusions. Interestingly, we found that the DOC domain of ZZEF1 binds directly to nucleosomes, independent of H3 histone modifications (**Fig. 6A** and **S6C**). These results suggest that ZZEF1 directly interacts with nucleosomes via its DOC domain, indicating that ZZEF1 may serve as a “pioneer” transcription factor for tRNA-Lys-TTT-3 loci.

**Figure 6.**
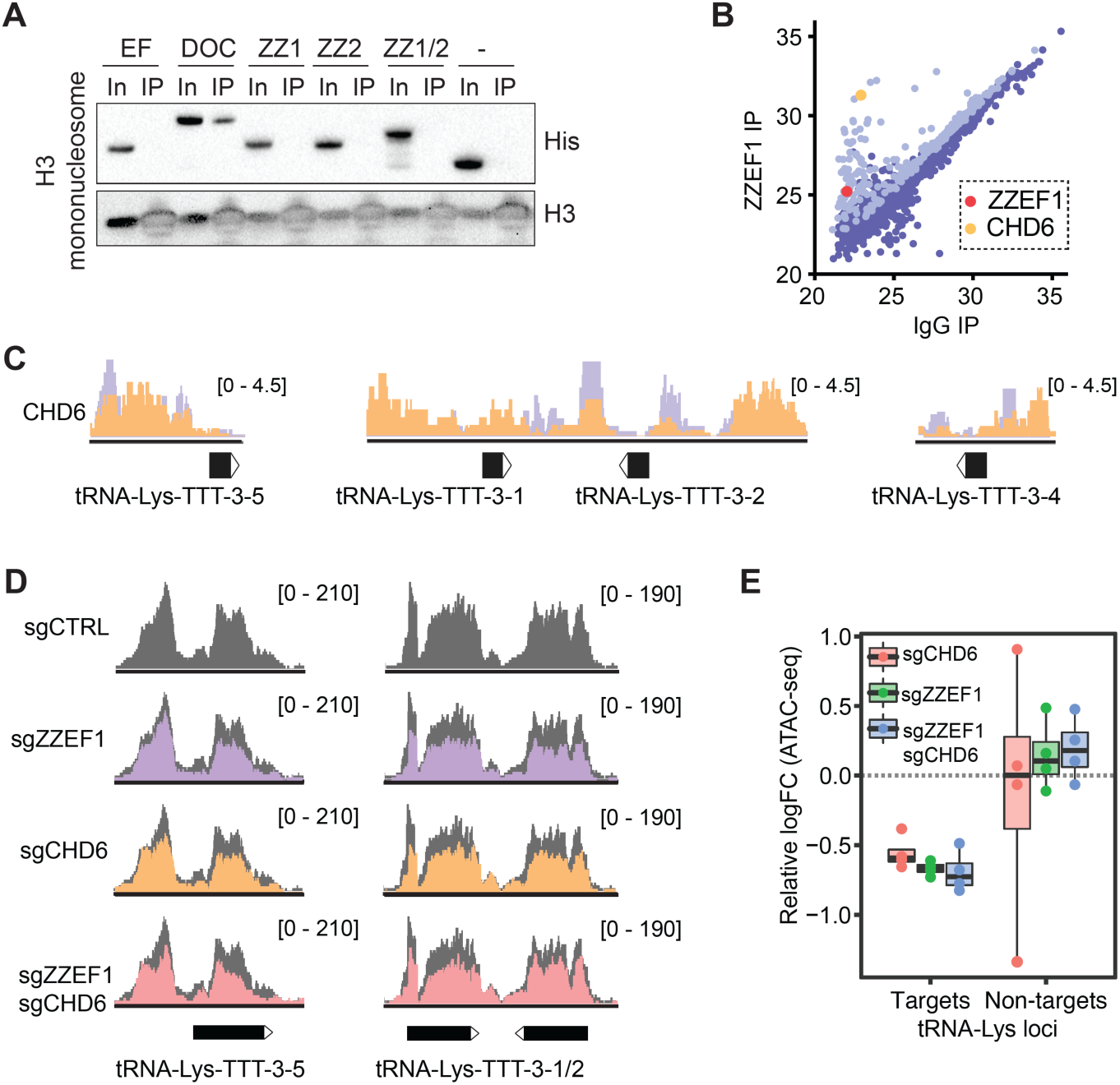
ZZEF1 cooperates with CHD6 to increase chromatin accessibility. **(A)** Pulldown immunoprecipitation (IP) results demonstrating interactions between ZZEF1 domains and mononucleosomes. In, input. IP, biotin pulldown. His, anti-His-tag western blot. **(B)** Dot plot showing CHD6 as a top interactor identified through ZZEF1 co-IP/MS. **(C)** Normalized ChIP-seq tracks showing enriched CHD6 binding (orange) in the proximal regions of tRNA-Lys-TTT-3 loci using a CHD6-specific antibody, compared to an IgG control in MDA cells. The purple tracks represent ZZEF1 binding from Fig. 5A. **(D)** Normalized ATAC-seq tracks showing chromatin accessibility at tRNA-Lys-TTT-3 loci in MDA cells with ZZEF1 knockdown, CHD6 knockdown, and CHD6 knockdown in a ZZEF1-deficient background, compared to control cells. **(E)** Boxplot showing statistical analysis of chromatin accessibility from ATAC-seq at the tRNA-Lys-TTT-3 loci and non-target loci in ZZEF1 knockdown, CHD6 knockdown, and CHD6 knockdown in a ZZEF1-deficient background compared to control cells.

Transcription factors typically cooperate with other factors to exert their regulatory effects. To determine the interactome of ZZEF1, we performed ZZEF1 co-immunoprecipitation coupled with mass spectrometry (co-IP/MS) experiments in MDA cells. Among the top interactors, we identified CHD6 (**Fig. 6B**), a known ATP-dependent chromatin remodeler^56^. This interaction was further validated by ZZEF1 co-IP followed by western blotting (**Fig. S6D**). Given that CHD6 functions in chromatin remodeling, and our observation of ZZEF1’s interaction with nucleosomes (**Fig. 6A** and **S6C**), we hypothesized that CHD6 collaborates with ZZEF1 to regulate chromatin accessibility and transcriptional activity at the tRNA-Lys-TTT-3 loci. To test this, we first performed ChIP-seq with a CHD6-specific antibody and compared the binding profiles to ZZEF1 ChIP-seq data. Interestingly, CHD6 binding significantly overlapped with ZZEF1 binding on a genome-wide scale (**Fig. S6E**, Jaccard index = 0.24, *P* = 0.0013, permutation test). Remarkably, CHD6 binding was highly enriched near the target tRNA-Lys-TTT-3 loci, closely mirroring ZZEF1’s binding pattern (**Fig. 6C** and **S6F-G**). These results suggest that CHD6 works in tandem with ZZEF1 to directly stimulate Pol III transcription of tRNA-Lys-TTT-3.

ATP-dependent chromatin remodelers regulate targets by repositioning nucleosomes to generate chromatin accessibility^57^. We hypothesized that ZZEF1 recruits CHD6 to generate chromatin accessibility at tRNA-Lys-TTT-3 loci. To test this, we conducted assays for transposase-accessible chromatin using sequencing (ATAC-seq) in MDA cells with ZZEF1 knockdown and CHD6 knockdown, respectively. Chromatin accessibility changes following CHD6 knockdown were significantly positively correlated with those observed after ZZEF1 knockdown (**Fig. S6H**, *R* = 0.475, *P* < 10^-100^, Pearson correlation), supporting a coordinated regulatory mechanism. Most importantly, we observed a significant reduction in chromatin accessibility around the target tRNA-Lys-TTT-3 loci following both ZZEF1 and CHD6 knockdown, while non-target tRNA-Lys loci remained unchanged (**Fig. 6D-E**). This suggests that ZZEF1 and CHD6 directly remodel chromatin at tRNA-Lys-TTT-3 loci. Additionally, we found that ZZEF1 binding potential—estimated by our ATAC-seq data—was decreased at these loci upon ZZEF1 knockdown (**Fig. S6I**), suggesting that ATAC-seq footprinting effectively captures ZZEF1 binding dynamics. We next applied this approach to TCGA-BRCA

ATAC-seq data^58^, consistently, we found that reduced binding potential at ZZEF1-specific sequence contexts was associated with decreased chromatin accessibility at tRNA-Lys-TTT-3-1 locus in breast cancer samples (**Fig. S6J**). Chromatin accessibility at these loci also positively correlated with accessibility in flanking regions of ZZEF1-specific sequence features (**Fig. S6J**). These results suggest that ZZEF1 acts as a locus-specific regulator of chromatin accessibility, specifically increasing accessibility around target loci to facilitate transcriptional activity.

To further investigate the epistatic relationship between ZZEF1 and CHD6 in chromatin remodeling at tRNA-Lys-TTT-3 loci, we also examined chromatin accessibility alterations using ATAC-seq in CHD6 knockdown cells in a ZZEF1-depleted background. We found that the double ZZEF1/CHD6 knockdown did not result in significantly different chromatin accessibility at tRNA-Lys-TTT-3 loci compared to the single knockdown cells (**Fig. 6D** and **6E**), further supporting the hypothesis that CHD6 and ZZEF1 collaborate to modulate chromatin accessibility and promote transcriptional activity at tRNA-Lys-TTT-3 loci. Additionally, a comparison of tRNA expression profiles in CHD6 knockdown MDA cells and CHD6 knockdown cells in a ZZEF1-depleted background revealed no significant differences in tRNA-Lys^UUU^ expression (**Fig. S6K**). Together, these findings indicate that CHD6 works in concert with ZZEF1 to regulate tRNA-Lys^UUU^ transcription by modulating chromatin accessibility at its target loci.

## Discussion

In this work, we present EXTRNA, the most comprehensive atlas (to our knowledge) to date of tRNA expression across human tissues. The dynamic expression patterns captured by EXTRNA, together with prior observations^20,24^, suggest that tRNAomes are subject to diverse regulatory modes. By leveraging an integrative approach involving EXTRNA, TCGA, CCLE, and a mixture of *in vitro* and *in vivo* experiments, we identified ZZEF1 as a previously uncharacterized transcriptional regulator that directly promotes the transcription of a specific tRNA, tRNA-Lys^UUU^. ZZEF1 interacts with the ATP-dependent chromatin remodeling enzyme CHD6 to facilitate Pol III transcription. Disruption of this regulatory axis impairs the translational efficiency of AAR codon-enriched proteins, including the tumor suppressor STK3, and promotes metastasis *in vivo*. Furthermore, clinical data link low ZZEF1 and STK3 expression to poorer survival outcomes in breast cancer, highlighting their clinical significance (**Fig. 7**). The EXTRNA dataset thus provides a valuable resource for deciphering the dynamic regulation of tRNA expression in cancer and offers a foundation for further mechanistic studies. Altogether, our study underscores the intricate regulation of tRNA expression and provides a framework for understanding tRNA-mediated molecular heterogeneity, with implications for novel therapeutic opportunities in cancer treatment.

**Figure 7.**
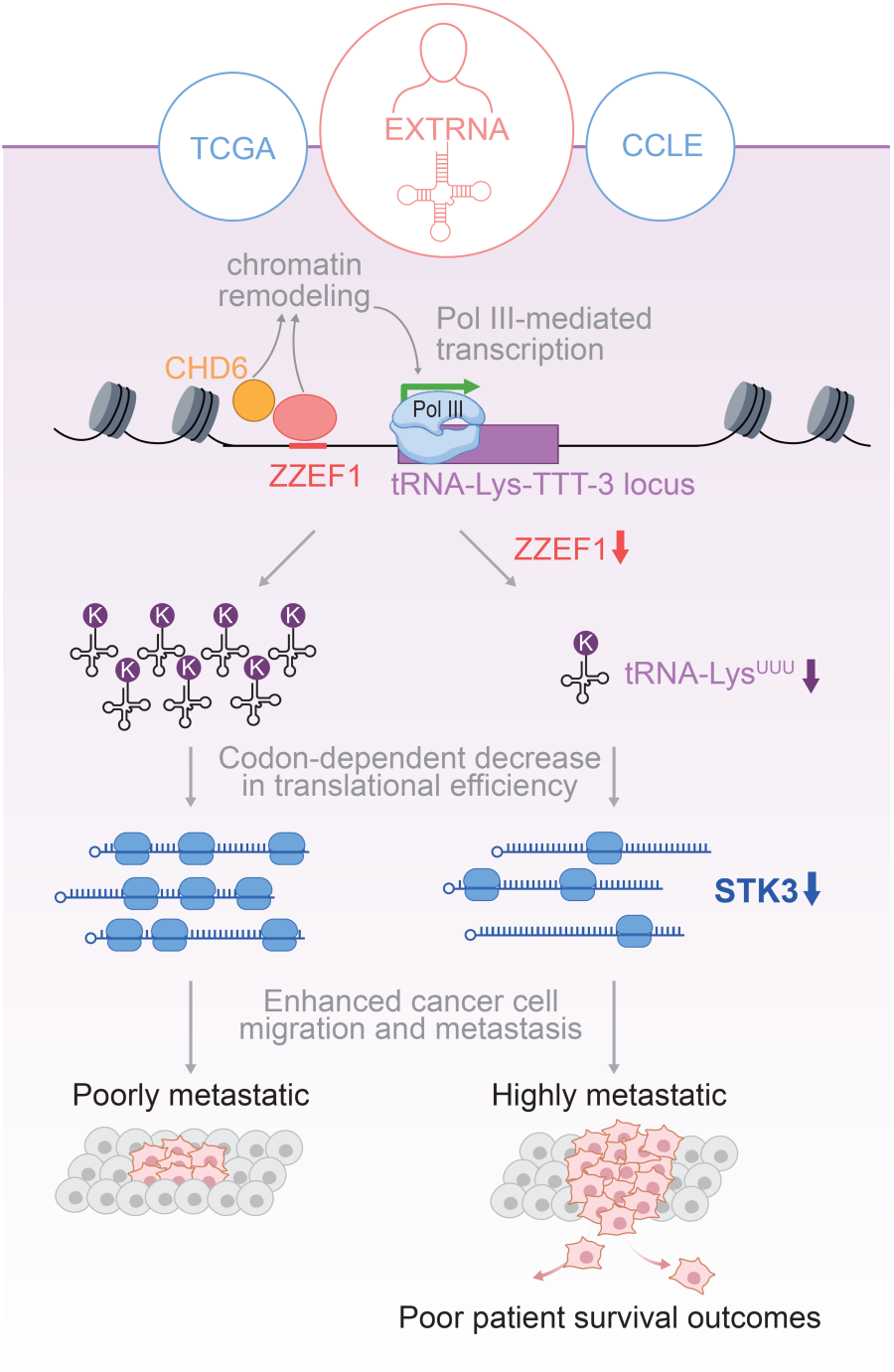
ZZEF1 promotes tRNA-Lys^UUU^ transcription through chromatin remodeling and drives codon-dependent effects on breast cancer metastasis. This schematic illustrates the regulatory mechanism of ZZEF1 revealed through integration of EXTRNA, TCGA, and CCLE datasets. ZZEF1 activates transcription of tRNA-Lys^UUU^-3 by engaging chromatin remodeling machinery, thereby enhancing the translational efficiency of AAR codon-enriched transcripts, including the tumor suppressor STK3. This codon-dependent mechanism contributes to the regulation of metastatic potential in breast cancer.

More broadly, our work raises fundamental scientific questions about the regulation of tRNA expression. Although RNA Pol III–the core protein complex responsible for tRNA transcription—is known to be homeostatically regulated (i.e. downstream of mTORC1), the mechanisms capable of sculpting tRNA expression have not previously been identified. Our findings support a model in which locus-specific factors, such as ZZEF1, selectively modulate Pol III activity, adding a previously underappreciated layer of regulatory complexity. This regulation is mediated through specific DNA sequence motifs, identified and characterized using ChIP-seq, a method well-suited for determining precise binding preferences. Supporting this, TCGA ATAC-seq data revealed consistent changes in chromatin accessibility associated with motif binding, while weaker signals at other loci were likely masked by methodological differences (**Fig. S6L**).

Moreover, our work highlights the importance of chromatin context in tRNA gene regulation. tRNA loci are known to be non-randomly positioned within nucleosomes^59^, but the underlying mechanisms and possible impacts remain largely unexplored. Previous studies observed that actively transcribed tRNA genes are often located near *cis*-regulatory elements, such as promoters or enhancers of Pol II-transcribed genes^37,38^, suggesting potential coordination between transcriptional programs. In our study, we demonstrate that loss of ZZEF1 binding reduces chromatin accessibility at the tRNA-Lys-TTT-3 loci, leading to decreased Pol III transcription and promotion of breast cancer metastasis via the ZZEF1–tRNA-Lys^UUU^–STK3 axis. While our model centers on tRNA-Lys^UUU^, the chromatin-level regulation we observed may extend to additional loci. Notably, our functional experiments show that perturbation of this axis alone is sufficient to drive metastasis (**Fig. 4**), establishing a direct link between chromatin accessibility, tRNA transcription, and tumor progression. Building on these findings, future studies will be essential to explore how chromatin architecture shapes Pol III activity within the broader transcriptional landscape—including whether regulators like ZZEF1 influence nearby Pol II genes, act beyond tRNA loci, or contribute to cross-talk between RNA polymerases.

Importantly, we demonstrate the physiological relevance of this regulatory mechanism in the context of breast cancer metastasis. Altered transcription of tRNA-Lys^UUU^ is sufficient to impair translation of AAR codon-enriched proteins, including STK3, thereby driving metastasis in vivo.

Subsequent analyses of clinical data link low ZZEF1 and STK3 expression to poorer survival outcomes in breast cancer, further highlighting the clinical and physiological significance of this discovery (**Figure 7**), underscoring both the biological and clinical impact of this regulatory axis. Our study provides mechanistic insight into how changes in tRNA expression contribute to cancer progression.

In sum, our findings reveal a previously unappreciated layer of tRNA gene regulation, define a new chromatin-based mechanism for Pol III transcriptional control, and highlight how deregulated chromatin remodeling factors, such as ZZEF1, can reprogram Pol III transcription and impact codon-specific translation to drive cancer development. Together, these insights provide a blueprint for decoding tRNA regulatory networks. They not only advance our understanding of non-coding RNA regulation but also point to new therapeutic strategies centered on the regulatory architecture of the tRNAome.

Looking forward, EXTRNA represents a valuable community resource. It enables comparative analyses of tRNA dynamics across tissues and conditions and lays the groundwork for identifying additional regulatory programs governing tRNA expression. We anticipate its broad application in uncovering tRNA-specific regulons across diverse biological processes and diseases, including cell-type specification and developmental transitions. Characterizing these regulons promises to advance in our understanding of how the central dogma of biology is regulated, while potentially uncovering new therapeutic targets.

## Acknowledgments

We thank Brian Plosky and Chiara Ricci-Tam for their suggestions and comments during manuscript preparation. We acknowledge the UCSF Center for Advanced Technology (CAT) for high throughput sequencing and other genomic analyses. We thank the Preclinical Therapeutics core as well as the Laboratory Animal Resource Center (LARC) at UCSF. We acknowledge support from our colleagues at the Helen Diller Family Comprehensive Cancer Center and the Breast Oncology Program. We thank Patricia Duchambon for her assistance in protein purification. H.G. is an Arc Core Investigator and research in this paper was partly funded by the Arc Institute. H.G. is an Era of Hope Scholar (W81XWH-2210121) and supported by grants from NCI (R01CA240984 and R01CA244634). S.F.T. was supported by grants from the National Cancer Institute of the National Institutes of Health under award numbers R35CA274446, U54CA261701, R01CA257153, the Hess Family foundation, the Breast Cancer Research Foundation, a Chan Zuckerberg Investigator award, and the Black Family Metastasis Center. V.R. acknowledges funding from NIH grant DP2-HG012442, the Searle Scholars Program, and the W.M. Keck Foundation. A.N. is a recipient of the DoD PRCRP Horizon Award W81XWH-19-1-0594, and is supported by ATIP-Avenir Program (CNRS/ARC Foundation), PSL Young Researcher Starting Grant, Prix Ruban Rose Avenir, and La Ligue Contre le Cancer. D.M. is supported by Boehringer Ingelheim Fonds MD fellowship. UCSF Laboratory for Cell Analysis shared resource facility is supported by NIH P30CA082103.

## Author contributions

H.G. and S.T. conceived the project. S.C., D.M., M.K. (Karimzadeh), Y.L., M.K. (Khoroshkin), B.B., S.L., C.C., P.N., K.G., T.J., C.M., B.H., H.M., V.R., S.T., A.N., and H.G. designed, performed, and analyzed the experiments and results. H.G., A.N., S.T., and V.R. acquired funding. S.C., A.N., M.K., H.G., and S.T. supervised the research. S.C., A.N., D.M., Y.L., and H.G. wrote the original draft. S.C., A.N., H.G., S.T., and V.R. reviewed and edited the manuscript.

## Data and materials availability

All sequencing data have been deposited in the GEO database under accession GSEXXX. The human breast cancer data were derived from the TCGA (at Genomic Data Commons, https://gdc.cancer.gov). The CPTAC breast cancer data set was obtained from Proteomics Data Commons (https://proteomic.datacommons.cancer.gov/pdc/). All other data supporting the findings of this study are available from the corresponding author on reasonable request.

## Supplementary Materials

### Materials and Methods

This study complies with all relevant ethical regulations that are approved by UCSF Institutional Review Board (IRB) and Institutional Animal Care and Use Committee (IACUC, approval number AN179718).

### Statistics and reproducibility

For *in vivo* experiments, mice were distributed into cohorts with 5 mice per cohort, which in NSG background is enough to observe a >2-fold difference with 90% confidence. For other experiments, no statistical methods were used to calculate sample size.

No data were excluded from the analyses.

Mice for *in vivo* experiments were randomly assigned into cohorts. For other experiments, no randomization was performed.

### Cell culture

All cells were cultured in a 37°C 5% CO2 humidified incubator. MDA-MB-231 (ATCC HTB-26), HEK293T (ATCC CRL-3216), A375 (ATCC CRL-1619), MCF7 (ATCC HTB-22), MIA PaCa-2 (ATCC CRL-1420), PANC-1 (ATCC CRL-1469), A2058 (ATCC CRL-3601) cell lines were cultured in DMEM high-glucose medium containing 1 mM sodium pyruvate. A549 (ATCC CCL-185), PC-3 (ATCC CRL-1435) cell lines were cultured in F12 medium. Hep G2 (ATCC HB-8065) cell line was cultured in EMEM medium supplemented with non-essential amino acids (NEAA). HCT 116 (ATCC CCL-247), SW480 (ATCC CCL-228) cell lines were cultured in McCoy’s 5A medium. MDA-MB-453 (ATCC HTB-131), SW620 (ATCC CCL-227) cell lines were cultured in Leibovitz’s L-15 medium.

HCC1806 (ATCC CRL-2335), C4-2B (ATCC CRL-3315), H23 (ATCC CRL-5800), K562 (ATCC CCL-243), LNCaP (ATCC CRL-1740), COLO 320 (ATCC CCL-220), ZR-75-1 (ATCCCRL-1500), BxPC-3 (ATCC CRL-1687), H1299 (ATCC CRL-5803), H358 (ATCC CRL-5807), HCC44 (DZMC ACC-534) cell lines were cultured in RPMI-1640 medium containing 25 mM HEPES. All growth media contained 2 mM L-glutamine and were supplemented with 10% FBS, penicillin (100 units/mL), streptomycin (100 μg/mL) and amphotericin B (1 μg/mL) (Gibco). All cell lines were routinely screened for mycoplasma with a PCR-based assay and tested negative.

### RNA isolation

Total RNA for RNA-seq and RT-qPCR was isolated using the Zymo QuickRNA isolation kit with in-column DNase treatment per the manufacturer’s protocol.

### Mature tRNA sequencing

Y-shaped Adapter MAture tRNA (YAMAT) sequencing was adapted from published study^1^ to include unique molecular identifiers (UMIs) in order to avoid overamplification of individual tRNA isodecoders. In total, 5 µg total RNA was deacetylated to remove amino acids from 3’ ends and demethylated to remove m^1^A, m^1^C, and m^3^C modifications using a proprietary demethylation mix (Arraystar, Inc). In total, 40 µm of YAMAT forked linkers were then incubated with pure demethylated and deacetylated RNA followed by the addition of 10× annealing buffer (50 mM Tris HCl pH 8, 100 mM MgCl_2_, 5 mM EDTA) and then overnight incubation with T4 RNA ligase 2.

Linker-ligated RNA was then incubated with RT Primer, and reverse transcription was performed with Superscript III RT (Invitrogen) followed by bead purification. Libraries were amplified using Phusion Hotstart II Polymerase (Thermo) with primers and indexes from the TruSeq Small RNA kit (Illumina). Amplified libraries were then bead-purified and run on the Agilent Bioanalyzer for confirmation of library size and quantification. The libraries were sequenced as PE75 runs on MiSeq sequencer (Illumina).

Raw sequencing reads were initially processed using UMI-tools^2^ to extract unique molecular identifiers (UMIs), followed by Cutadapt with parameters -e 0.12 -m 20 to trim adapter sequences^3^. Deduplication was then performed based on the extracted UMIs. Paired-end reads were merged into fragments using PEAR^4^ and aligned to a human tRNA reference using Bowtie2^5^ with the --very- sensitive option. Fragment counts for each tRNA gene were aggregated at the isodecoder level and normalized using the varianceStabilizingTransformation function in DESeq2^6^. For isoacceptor-level expression, fragment counts from corresponding isodecoders were summed and subsequently normalized using the same DESeq2 transformation.

### MOFA+ joint analyses of tRNA expression and mRNA expression

tRNA isoacceptor expression profiles (from our dataset) and mRNA expression profiles (from CCLE)^7^ were extracted for overlapping cell lines. Gene function annotations were downloaded from the Human MSigDB Collections^8^ using the file c5.go.v2024.1.Hs.symbols.gmt. Genes associated with transcriptional regulation, chromatin remodeling, or histone binding—defined by gene ontology terms containing HISTONE_BINDING, CHROMATIN_REMODELING, or

REGULATION_OF_DNA_TEMPLATED_TRANSCRIPTION—were selected for downstream analysis.

Joint analysis was performed using MOFA+^9^, with the number of latent factors set to 16. Both mRNA and tRNA modalities were modeled using Gaussian likelihoods. The training procedure was initialized with a fixed random seed (42) to ensure reproducibility, and the model was trained in “fast” convergence mode. Evidence Lower Bound (ELBO) estimation began after the first iteration and was evaluated at every iteration. The maximum number of training iterations was set to 1,000 to ensure model convergence.

For each latent factor identified by MOFA+, we extracted feature weights from both RNA and tRNA modalities and retained those with absolute values greater than 0.1. *K*-means clustering was then applied to group the selected features. To explore potential regulatory modules, we constructed RNA– tRNA co-expression networks within each cluster. Prior to network construction, we confirmed co-expression by calculating Spearman correlations across cell lines, retaining only feature pairs with an absolute correlation coefficient greater than 0.25 as edges in the network.

### Master regulator inference from TCGA-BRCA dataset

Matched mRNA and small RNA expression data were obtained from the TCGA-BRCA cohort^10–12^. Expression values were first winsorized and rank-transformed. We applied the ARACNe algorithm to infer mutual information–based dependencies between mRNAs and tRFs across samples^13,14^. Significant interactions (FDR < 0.01) were determined by permutation testing, and indirect associations were removed using the data processing inequality. The resulting mRNA–tRF network was used for master regulator analysis based on a bimodality scoring approach. Candidate regulators were further filtered by intersecting with nucleic acid-binding protein and nuclease gene sets.

### Gene knockdown and overexpression

MDA-MB-231 and HCC1806-LM2 cells expressing dCas9-KRAB fusion protein were constructed by lentiviral delivery of pMH0006 (Addgene #135448) and FACS isolation of BFP-positive cells.

The lentiviral constructs were co-transfected with pCMV-dR8.91 and pMD2.D plasmids using TransIT-Lenti (Mirus) into 293T cells, following manufacturer’s protocol. Virus was harvested 48 hours post-transfection and passed through a 0.45 µm filter. Target cells were then transduced overnight with the filtered virus in the presence of 8 µg/mL polybrene (Millipore).

Guide RNA sequences for CRISPRi-mediated gene knockdown were cloned into pCRISPRia-v2 (Addgene #84832) via BstXI-BlpI sites. For double knockdown experiments, pCRISPRia-v2 plasmid was modified to construct pCRISPRia-v2-Blast, replacing puromycin acetyltransferase by blasticidin deaminase coding sequences. After transduction with sgRNA lentivirus, the cells were selected with 1.5 µg/mL puromycin or 20 µg/mL blasticidin (Gibco). Knockdown of target genes was assessed by RT-qPCR as described below.

tRNA-Lys^UUU^-3 isodecoder was cloned into pLKO.1 vector for overexpression. After transduction, the cells were selected with 20 µg/mL blasticidin.

### RT-qPCR

Transcript levels were measured using quantitative RT-PCR by reverse transcribing total RNA to cDNA (Maxima H Minus RT, Thermo), then using PerfeCTa SYBR Green SuperMix (QuantaBio) per the manufacturer’s instructions. HPRT1 was used as endogenous control. For tRNA experiment, reverse transcription was performed with Superscript III RT (Invitrogen), and 5S rRNA was used as endogenous control in RT-qPCR.

### Small RNA sequencing

Small RNA-seq libraries were constructed using NextFlex Small RNA Library Kit V3 (BIOO Scientific), with 100 ng small RNA fraction as input, according to manufacturer’s recommendations. The libraries were sequenced as SE50 runs on Illumina HiSeq4000 instrument at UCSF Center for Advanced Technologies.

To process the data, fastq files were first trimmed using cutadapt (v2.2). UMIs were then extracted using umi_tools (v1.0.0), aligned with bowtie2 (2.3.5) with --sensitive --end-to-end -N 1 flags. Aligned reads were sorted using samtools (v1.9) and deduplicated. Small RNAs were then counted using bedtools (v2.28) against an annotated feature space of small RNAs. The resulting count matrices were then compared using DESeq2 (v1.24).

### Western Blotting

Cell lysates were prepared by lysing cells in ice-cold RIPA buffer (25 mM Tris-HCl pH 7.6, 0.15 M NaCl, 1% IGEPAL CA-630, 1% sodium deoxycholate, 0.1% SDS) containing 1× protease inhibitors (Thermo Scientific). Lysate was cleared by centrifugation at 20,000 × g for 10 min at 4°C. Samples were denatured for 10 min at 70°C in 1× LDS loading buffer (Invitrogen) and 50 mM DTT. Proteins were separated by SDS-PAGE using 4-12% Bis-Tris NuPAGE gels, transferred to nitrocellulose (Millipore), blocked using 5% BSA in PBST, and probed using target-specific antibodies. Bound antibodies were detected using HRP-conjugated secondary antibodies (anti-mouse IgG, Light chain specific, HRP (Cell Signaling 91196S, 1/5,000) or anti-rabbit IgG, conformation specific, HRP (Cell Signaling 5127S, 1/5,000)) and ECL substrate (Pierce) or infrared dye-conjugated secondary antibodies (anti-mouse IgG, IRDye 800CW (Li-Cor 926-32210, 1/10,000) or anti-rabbit IgG, IRDye 680RD (Li-Cor 926-68071, 1/10,000)) according to the manufacturer’s instructions.

Primary antibodies: beta-tubulin (Proteintech 66240-1-Ig, 1/10,000), GAPDH (Proteintech 60004-1-Ig, 1/10,000), STK3 (Proteintech 12097-1-AP, 1/1,000), ZZEF1 (Bethyl, A304-068A, 1/2,000), CHD6 (Bethyl, A301-221A, 1/1,000).

### Northern blotting

Total RNA was separated on 10% TBE-urea gels at 250 V for 1.5-2 h and transferred to Hybond-N+ membranes for 1 h. RNA was cross-linked to the membrane at 240 mJ/cm², then pre-hybridized with UltraHyb-Oligo buffer at 42°C for 1 h. Oligonucleotide probes were labeled with [³²P]ATP using T4 PNK (NEB), purified with G25 columns, and hybridized overnight at 42°C. Blots were washed twice and exposed to phosphor screens. Band intensity was quantified using ImageJ and normalized to U6 abundance.

### Ribosome profiling

Ribosome profiling was performed as previously described^15^ with following modifications. The RNA concentration in the lysates was determined with the Qubit RNA HS kit (Thermo). Monosomes were isolated using MicroSpin S-400 HR (Cytiva) columns, pre-equilibrated with 3 mL polysome buffer per column. 100 µl digested lysate was loaded per column (two columns were used per 200 µl sample) and centrifuged 2 min at 600 × g. The RNA from the flow through was isolated using the Zymo RNA Clean and Concentrator-25 kit. In parallel, total RNA from undigested lysates were isolated using the same kit.

Libraries were sequenced on a SE50 run on Illumina HiSeq4000 instrument at UCSF Center for Advanced Technologies.

To process the reads, the Ribo-seq reads were first trimmed using cutadapt (v2.3) to remove the linker sequence AGATCGGAAGAGCAC. The fastx_barcode_splitter script from the Fastx toolkit (v0.0.13) was then used to split the samples based on their barcodes. Since the reads contain unique molecular identifiers (UMIs), they were collapsed to retain only unique reads. The UMIs were then removed from the beginning and end of each read (2 and 5 Ns, respectively) and appended to the name of each read. Bowtie2 (v2.3.5) was then used to remove reads that map to ribosomal RNAs and tRNAs, and the remainder of reads were then aligned to mRNAs (we used the isoform with the longest coding sequence for each gene as the representative). Subsequent to alignment, umitools (v0.3.3) was used to deduplicate reads.

### RNA sequencing (RNA-seq)

RNA-seq libraries (used for calculating translation efficiencies) were prepared using SMARTer Stranded Total RNA-Seq Kit v2 - Pico Input Mammalian (Takara), with 50 ng total RNA as input, according to manufacturer’s instructions. The libraries were sequenced as SE50 runs on Illumina HiSeq4000 instrument at UCSF Center for Advanced Technologies.

### Translational efficiency and codon usage modeling

Gene-level translational efficiency (logTER) was estimated using the ribolog suite, and codon-specific ribosome occupancy was inferred using CELP bias coefficients^16^. To assess the impact of the ZZEF1–tRNA-Lys^UUU^ regulatory axis on translation, we quantified the differential effect as the difference in logTER between tRNA-Lys^UUU^ overexpression in ZZEF1 knockdown cells and ZZEF1 knockdown cells. To evaluate codon usage contributions to translational efficiency changes, we fit a generalized linear model with logTER as the response variable and codon usage frequencies as predictors. Codons with significant coefficients (*P* < 0.05) were considered to have a statistically measurable influence on translational efficiency.

### Chromatin immunoprecipitation and sequencing (ChIP-seq)

MDA-MB-231 cells (10×10^6^ per replicate) were washed with ice-cold 1× PBS and fixed in 1% PFA in 1× PBS for 10 min at room temperature. The fixation was quenched with 100 mM glycine (final concentration) for 5 min. The cells were lysed in 100 µl lysis buffer (50 mM Tris HCl pH 8, 10 mM EDTA, 1% SDS) containing 1× protease inhibitors (Thermo Scientific) on ice for 10 min. The lysates were then diluted with 8 volumes of dilution buffer (20 mM Tris HCl pH 8, 150 mM NaCl, 1% Triton X-100, 2 mM EDTA, 3 mM CaCl_2_) with protease inhibitors and prewarmed to 37°C. The lysates were digested with micrococcal nuclease (MNase, Sigma N3755) at 1 U/mL final concentration, 2 min at 37°C, and the reaction was stopped by adding 10 mM EDTA and 20 mM EGTA (final concentrations). The lysates were sonicated 10 min using Bioruptor sonicator (Diagenode), at 30 sec on/off HIGH setting, and clarified 10 min at 21,000 g at 4°C.

The DNA was extracted from 50 µl chromatin preparation using 2× Elution buffer (100 mM NaHCO_3_, 350 mM NaCl, 0.8% SDS) supplemented with 0.2 µg/mL (final concentration) RNase A (Thermo) and incubating at 65°C overnight. The reactions were supplemented with 0.4 µg/mL (final concentration) Proteinase K (Thermo), incubated 1h at 60°C, and the DNA was purified using DNA Clean and Concentrator 25 kit (Zymo). The DNA was analyzed by agarose gel electrophoresis, and chromatin preparations were used for IP if the obtained fragments were 100-500 bp in length.

900 µL chromatin preparation was used for IP with 5 µg anti-ZZEF1 (Bethyl, A304-068A) or anti-CHD6 (Bethyl, A301-221A) at 4°C overnight. In parallel, 100 µl chromatin was processed as input. The next day, the samples were clarified 5 min at 21,000 g at 4°C, and the supernatants were added to Protein A/G magnetic beads (Pierce). After 4 h at 4°C, the beads were washed once in WB1 (20 mM Tris HCl pH 8, 150 mM NaCl, 2 mM EDTA, 1% Triton X-100, 0.1% SDS), three times in WB2 (20 mM Tris HCl pH 8, 500 mM NaCl, 2 mM EDTA, 1% Triton X-100, 0.1% SDS), once in WB3 (10 mM Tris HCl pH 8, 250 mM LiCl, 1 mM EDTA, 1% sodium deoxycholate, 1% Igepal CA-630), and twice in TE (10 mM Tris HCl pH 8, 1 mM EDTA). The DNA was eluted in elution buffer (20 mM Tris HCl pH 8, 300 mM NaCl, 10 mM EDTA, 5 mM EGTA, 1% SDS) supplemented with 0.25 µg/mL Proteinase K (Thermo), at 65°C overnight. The DNA was extracted once with phenol-chloroform-isoamyl alcohol, once with chloroform, ethanol precipitated, and dissolved in 10 µL 10 mM Tris HCl pH 8.

ChIP DNA or 100 ng of input DNA was used for library preparation using NEBNext Ultra II DNAseq Kit (NEB E7103S). The libraries were sequenced as PE35 runs on NextSeq 550 sequencer (Illumina). To process the data, we used the ENCODE3 v1 pipeline. Briefly stated, reads were aligned using bowtie2 (v2.3.5), deduplicated using Picard (v2.21.3), and converted to BED files using bedtools (v2.28.0). UNIX shuf command was used to split into random psuedoreplicates, and reads were filtered against the ENCODE blacklist. Finally, MACS2 (v2.2.7) was used to call peaks. Reads mapping to the relevant tRNA loci were used to compare samples.

### ZZEF1 binding motif discovery

We used DREME (v5.5.5)^17^ to identify ZZEF1 sequence motifs by using -maxk 20 and the default parameters on a ChIP-seq narrowPeak file of ZZEF1 binding peaks.

### Co-occupancy analysis of CHD6 and ZZEF1 binding sites

To assess the genomic co-localization of CHD6 and ZZEF1 binding, we first calculated the distances between significant ZZEF1 peaks and CHD6 peaks (obtained from our ChIP-seq data) using the calcFeatureDist function from the GenomicDistributions package^18^. The Jaccard index was computed by quantifying the proportion of overlapping 300K nucleotide peak regions between ZZEF1 and CHD6 ChIP-seq peaks. To evaluate the statistical significance of the observed overlap, a permutation test was performed. Specifically, 10,000 sets of randomly shuffled genomic regions— matched in length and number to the CHD6 peaks—were generated, and their Jaccard indices with ZZEF1 peaks were computed. The empirical *P* value was calculated as the fraction of permuted Jaccard indices greater than or equal to the observed value.

### Assay for transposase-accessible chromatin using sequencing (ATAC-seq)

The libraries were constructed using Fast-ATAC protocol^19^. Briefly, 10^5^ cells were resuspended in 50 µL Tn5 master mix and incubated 30 min at 36°C with mixing at 1000 rpm. The reaction was stopped by adding 100 µL DNA Binding buffer from DNA Clean and Concentrator 5 kit (Zymo). The DNA was purified using the kit and eluted in 20 µL elution buffer. Ten µL of the tagmented DNA was used for library amplification with NEBNext Ultra II 2× master mix (NEB) and Nextera P5/P7 index primers (Illumina) by 8 PCR cycles. The bead-purified libraries were sequenced as a PE35 run on NextSeq 550 sequencer (Illumina). Reads were then processed using the ENCODE ATAC-seq pipeline. Filtered narrowPeaks were merged across samples, and peaks overlapping tRNA loci were marked. Reads mapping each peak was then counted and compared between samples using DESeq2.

### Co-immunoprecipitation and mass spectrometry (CoIP-MS)

MDA-MB-231 cells were washed three times with ice-cold PBS and lysed in lysis buffer (20 mM Tris-HCl pH 7.4, 150 mM NaCl, 0.5% NP-40, 2 mM EDTA, protease inhibitor cocktail, and RNase inhibitor). Lysates were homogenized by passing through a 23-gauge needle and rotated at 4°C for 1–2 h. After centrifugation at 600 g for 20 min at 4°C, supernatants were collected. Sepharose beads were pre-washed in lysis buffer and incubated with lysates for preclearing. 450 µl of precleared lysate was incubated with 5–10 µg of anti-ZZEF1 antibody (or control IgG) at 4°C overnight with rotation. The next day, samples were incubated with pre-washed sepharose beads for 1 h at 4°C, washed four times with lysis buffer, and proteins were eluted for downstream analysis. Eluates were processed for mass spectrometry. LC-MS/MS was performed at the Rockefeller Proteomics Resource Center as previously described^20,21^.

### Co-immunoprecipitation and western blotting (CoIP-WB)

MDA-MB-231 cells (10×10^6^ per replicate) were washed with ice-cold 1× PBS and lysed in nuclei lysis buffer (50 mM Tris HCl pH 7.5, 1% SDS, 1 mM EDTA) containing 1× protease inhibitors (Thermo Scientific) on ice for 10 min. The lysates were then diluted with 8 volumes of dilution buffer (50 mM Tris HCl pH 7.5, 150 mM NaCl, 1% Triton X-100, 1 mM EDTA, 3 mM CaCl_2_) with protease inhibitors and prewarmed to 37°C. The lysates were digested with micrococcal nuclease (MNase, Sigma N3755) at 1 U/mL final concentration, 5 min at 37°C, and the reaction was stopped by adding 10 mM EDTA and 20 mM EGTA (final concentrations). The lysates were cleared 10 min at 21,000 g at 4°C and used for IP.

4 µg anti-ZZEF1 (Bethyl, A304-068A) or anti-CHD6 (Bethyl, A301-221A) antibody or rabbit IgG was used per IP, at +4°C overnight. The next day the IP reactions were clarified 5 min at 21,000 g at +4°C, and the supernatants were added to Protein A/G magnetic beads (Pierce). After 2h incubation at +4°C, the beads were washed three times in wash buffer (25 mM Tris HCl pH7.5, 150 mM NaCl, 0.05% Triton X-100) and the proteins were eluted in 1× NuPage LDS loading buffer (Thermo) with 100 mM DTT, 10 min at 70°C. The protein samples were separated onto 3-8% Tris-Acetate PAGE gels and analyzed by western blotting as described above.

### Recombinant ZZEF1 domain purification

Human ZZEF1 domain information was obtained from UniProt^22^*. E. coli* codon-optimized DNA sequences coding ZZEF1 domains (EF hand (111-146 amino acid), DOC (226-405 amino acid), ZZ1 (1778-1833 amino acid), ZZ2 (1827-1882 amino acid), ZZ1/2 (1778-1882 amino acid)) were synthesized and cloned into a His-MBP fusion expression vector (Addgene #29654) via BamHI site and Gibson assembly. The correct cloning was confirmed by Sanger sequencing. The expression vectors were then used to transform BL21 Star (DE3) cells (Thermo). To induce recombinant protein expression, overnight *E. coli* cultures were diluted 100× in LB medium supplemented with antibiotics and 0.05 mM ZnCl_2_, grown at 37°C until mid-exponential phase (OD_600_ = 0.5), and then switched to 18°C for overnight induction with 0.2 mM IPTG. Frozen bacterial pellets were resuspended in Lysis buffer (20 mM Tris pH 7.0, 250 mM NaCl, 10% glycerol, 10 mM imidazole, 5 mM TCEP, protease inhibitors) and lysed in French press at 2 kbar. The lysate was clarified at 39,000 g for 1 hour at 4°C, then loaded on a 5 mL HisTrap (Cytiva) column (equilibrated in Lysis buffer) using the ÄKTA (Cytiva) HPLC system. The column was washed with 15 column volumes (CV) of Wash buffer (20 mM Tris pH 7, 250 mM NaCl, 10% glycerol, 20 mM imidazole, 5 mM TCEP). The proteins were eluted in a linear gradient of 5 CV 20-400 mM imidazole in Elution buffer (20 mM Tris pH 7, 250 mM NaCl, 10% glycerol, 5 mM TCEP). The target protein containing fractions were pooled and dialysed in Storage buffer (20 mM Tris pH 7, 500 mM NaCl, 10% glycerol, 5 mM TCEP), in a 12-14 MWCO membrane. The protein concentration was determined with BCA assay (Pierce).

### Nucleosome binding assay

Recombinant mononucleosomes assembled with biotinylated DNA, containing unmodified H3.2, or H3K27me3, or H3K4me3/H3K27ac histones (Active Motif) were used for binding assays. Briefly, 1 µg of nucleosomes were incubated with 5 µg of recombinant ZZEF1 domain proteins in 1× binding buffer (10 mM Tris pH 7.5, 50 mM NaCl, 0.1 mM DTT, 0.05% Igepal CA-630), in a final volume of 100 µl, 30 mins on ice. 10% of the reaction was set aside as input and the remaining volume was used for streptavidin pull-down using streptavidin magnetic beads (NEB) washed in 1× Binding Buffer. After 3h incubation at 4°C, the beads were washed 4× in cold 1× Binding Buffer, and protein eluted in 1× Novex LDS sample buffer (Thermo) 10 mins at 70°C. The input and pull-down fractions were then analyzed by western blotting as described above, using anti-His tag-HRP antibody (Proteintech, #HRP-66005, 1/5000 dilution) or anti-H3 antibody (Diagenode, #C15200011, 1/10000 dilution).

### ZZEF1 protein structure prediction and illustration

ZZEF1 protein structure was predicted by AlphaFold3^23^. Domains were colored with PyMOL.

### ATAC-seq–based estimation of ZZEF1 binding potential and chromatin accessibility

For ATAC-seq data from sgCTRL and sgZZEF1 cells, reads from biological replicates were combined, and Tn5 insertions were quantified using bedtools by intersecting motif and peak regions with BAM files that had been Tn5-shifted using alignmentSieve --ATACshift. Insertion counts were separately extracted for motif binding sites (10 bp) and their flanking regions (±100 bp). For TCGA-BRCA, normalized bigWig files^24^ were used to quantify Tn5 insertion signals at motif sites, flanking regions, and tRNA loci using bigWigAverageOverBed. Flanking accessibility was calculated as the log₂-transformed normalized insertion signal in the flanking region. Footprint depth was defined as the log₂ ratio of insertion at the motif site over the flanking region, following previously established methods^24^.

### tRNA-Lys-TTT-3-pre-tRNA quantification

The relative expression of tRNA-Lys-TTT-3-pre-tRNA was measured by RT-qPCR by reverse transcribing total RNA to cDNA (Superscript III RT), then using PerfeCTa SYBR Green SuperMix (QuantaBio) per the manufacturer’s instructions. 5S rRNA as the endogenous control. Primer design followed previously published guidelines^25^, with the forward primer spanning the 5′ leader sequence and the first six bases of the tRNA, and the reverse primer spanning the last six bases of the tRNA and the 3′ trailer sequence. Expression values represent the average of two biological replicates, each with four technical replicates.

### Metastatic colonization assay

Mice were housed in accordance with UCSF IACUC protocol (approval number AN179718) in humidity- and temperature-controlled rooms on a 12 hour light-dark cycle with free access to food and water. 7-12 week-old age-matched female NOD *scid* gamma mice (NSG, Jackson Labs, 005557) were used for lung colonization assays. For this assay, cancer cells constitutively expressing luciferase were suspended in 100 μL PBS and then injected via tail-vein (5×10^4^ for MDA-MB-231, 1×10^5^ for HCC1806-LM2). Cancer cell growth was monitored *in vivo* at the indicated times by retro-orbital injection of 100 µl of 15 mg/mL luciferin (Perkin Elmer) dissolved in 1× PBS, and then measuring the resulting bioluminescence with an IVIS instrument and Living Image software (Perkin Elmer). Mice were euthanized before the normalized photon flux in the lung region reached 5×10^8^; this limit was not exceeded.

### Migration assay

MDA-MB-231 cells (sgCTRL, sgZZEF1, and sgSTK3) were serum-starved for 24 hours in medium lacking FBS prior to the migration assay. Transwell migration assays were conducted using 24-well plates with 8-μm pore-size filter inserts (Fisher Scientific, 0877121). 5×10⁴ cells were resuspended in serum-free medium and seeded into the upper chambers of the inserts in four biological replicates per sample. The lower chambers were filled with 600 μL of complete growth medium supplemented with 10% FBS (VWR, 97068086) as a chemoattractant. After 24 hours of incubation at 37 °C, non-migrated cells on the upper membrane surface were removed using a cotton swab. Cells that had migrated to the underside of the membrane were fixed with methanol (Millipore Sigma, 34860-1L-R) and stained with Crystal Violet (Millipore Sigma, HT90132-1L). Membranes were imaged, and migrated cells were quantified using a custom image analysis script.

#### Cell proliferation assay

Cell proliferation was measured using the CellTiter-Glo Luminescent Cell Viability Assay (Promega, G7570) in 96-well tissue culture plates (Thermo Scientific, 165306). At Day 0, MDA-MB-231 Cells (sgCTRL, sgZZEF1, and sgSTK3) were seeded at 1,000 cells per well in 100 μL of medium, in eight biological replicates per sample. Control wells containing only media were included to assess background luminescence. Three identical plates were prepared to measure cell growth on Day 1, Day 3, and Day 5. Background-subtracted values were used to estimate growth rates by fitting an exponential model, ln(*N_t_*_-1_/*N*_1_) = *rt*, *t* is the time in days and *r* is the proliferation rate. Two-sided *t*-tests were used to assess statistically significant differences between samples.

## Supplementary Figures

**Figure S1.**
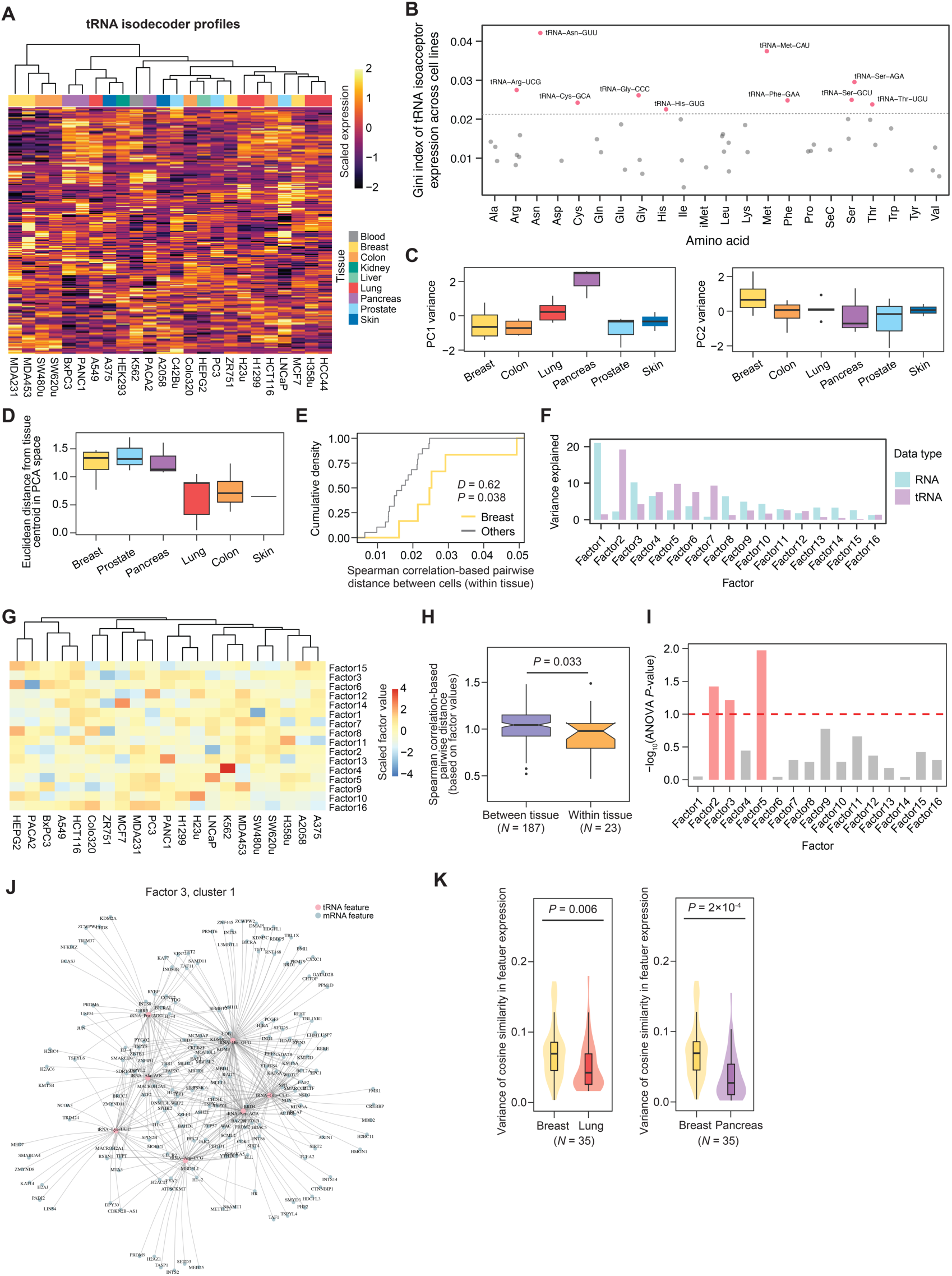
tRNA expression profiles in EXTRNA. **(A)** Heatmap illustrating tRNA isodecoder expression profiles in EXTRNA, with hierarchical clustering applied to group similar profiles. **(B)** Scatter plot showing the gini index distribution of tRNA isoacceptor expression across cell lines, representing variability in expression levels. tRNA isoacceptors with the highest variance (top 10 gini index values) are colored in red and labeled. **(C)** Boxplots showing the distributions of PC1 (left) and PC2 (right) values for cells from each tissue, as presented in Fig. 1D (only tissues with ≥2 cell lines are included). Association between PC values and tissue type was assessed using MANOVA (*P* = 0.04). **(D)** Boxplots showing the distribution of Euclidean distances from the tissue centroid in PCA space (from Fig. 1D) for each tissue (only tissues with ≥2 cell lines are included). Differences across tissues were evaluated using the Kruskal–Wallis test (*P* = 0.046). **(E)** Empirical cumulative distribution curves showing spearman correlation–based pairwise distances between cell lines within each tissue. Breast tissues were compared to other tissues in Fig. 1E. *D*-value and *P*-value were calculated using the Kolmogorov–Smirnov test. **(F)** Barplot showing the variance explained by each latent factor for tRNA and RNA datasets in the joint analysis. **(G)** Heatmap illustrating latent factor values across cell lines obtained from the joint analysis, with hierarchical clustering applied to group similar profiles. **(H)** Boxplot comparing Spearman correlation-based cell-cell pairwise distances calculated from **Fig. S1G** between tissue types and within tissue types. *P*-value was calculated using the Mann–Whitney *U* test. **(I)** Barplots showing the statistical significance of tissue-type associations for each latent factor, represented as -log_10_(*P*) values from ANOVA tests. Bars in red indicate latent factors with *P* < 0.1. (**J)** Regulatory network module (factor 3, cluster 1) identified from the joint analysis, similar to Fig. 1G but in a different module. **(K)** Violin and boxplot representation of the variance in expression similarity of tRNA features and RNA features within each module across cells from each tissue type. A few variance values out of the *y* axis range are not shown. *P* values were calculated using the Mann–Whitney *U* tests.

**Figure S2.**
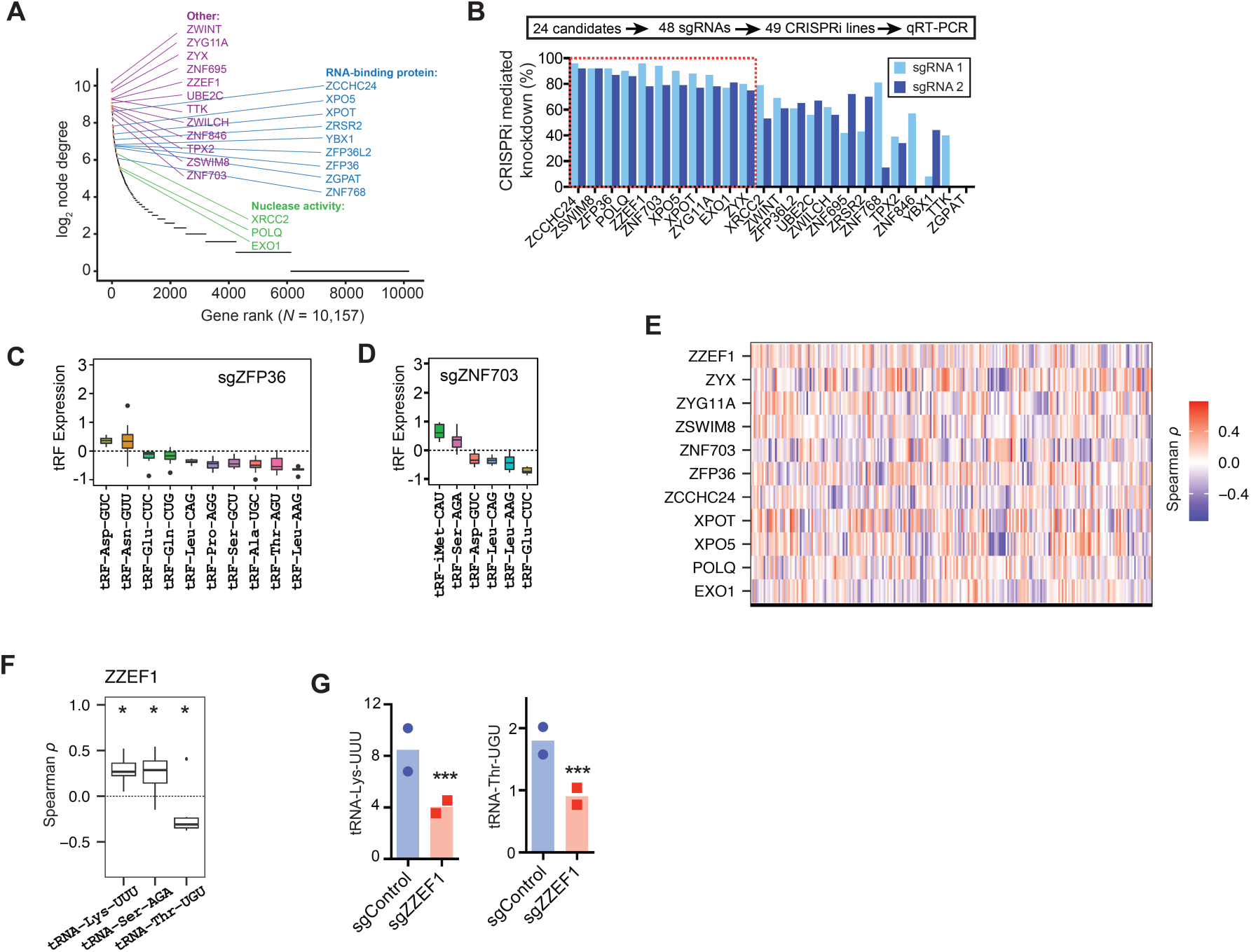
Validation of regulator candidates on tRF and tRNA expression. **(A)** Node degree-based ranking of identified transcriptional regulator candidates. Genes with nucleic acid binding capacity and/or nuclease activity were labeled. **(B)** Barplot showing CRISPRi knockdown efficiencies of 24 inferred regulator candidates in MDA cells. **(C)** Alterations in tRF expression in MDA cells with ZFP36 knockdown compared to control cells. **(D)** Similar to **(C)**, but showing results for ZNF703 knockdown. **(E)** Heatmap illustrating the expression correlation between individual tRNA isodecoders (EXTRNA) and regulator candidates (CCLE) across the cancer cell lines. **(F)** Boxplots depicting significant co-expression patterns (*P* < 0.05) between specific tRNAs and ZZEF1. *, *P* < 0.05. *P*-values for the association between tRNA isoacceptors and ZZEF1 were obtained by combining *P*-values from Spearman correlation analyses between tRNA isodecoder expression and ZZEF1 using Fisher’s combined *P*-value method. (**G)** Barplot showing mature tRNA-Lys^UUU^ and tRNA-Thr^UGU^ expression levels in MDA cells with ZZEF1 knockdown compared to control cells.

**Figure S3.**
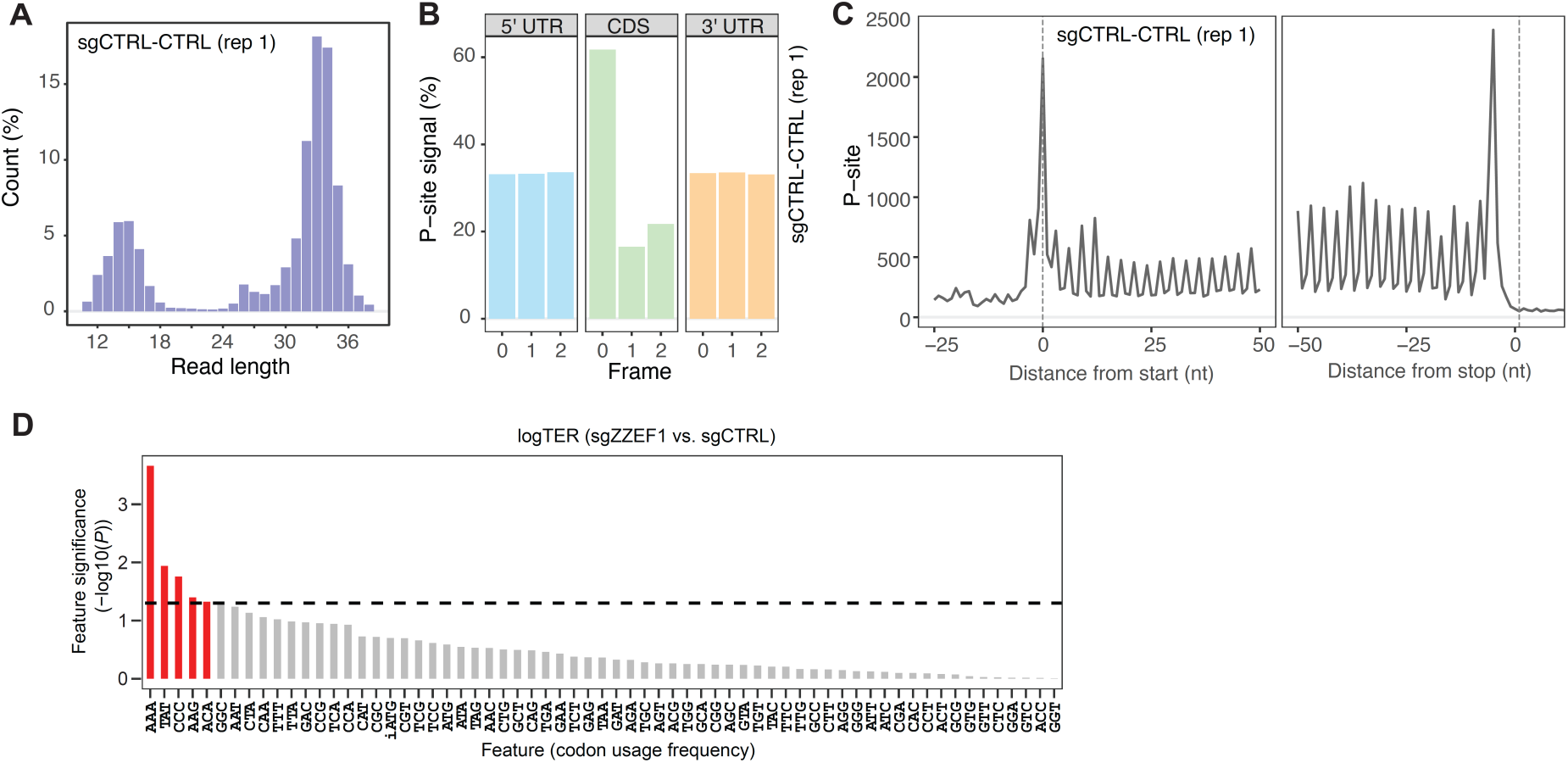
Ribo-seq profiling captures translational dynamics. **(A)** Read length distribution of ribosome profiling (Ribo-seq) data. **(B)** Three-nucleotide periodicity observed in coding regions and untranslated regions (UTRs). **(C)** Read count distribution around start and stop codons in Ribo-seq data. **(D)** Bar plot showing codon usage feature importance in accounting for translational efficiency alterations in MDA cells after ZZEF1 knockdown compared to controls.

**Figure S4.**
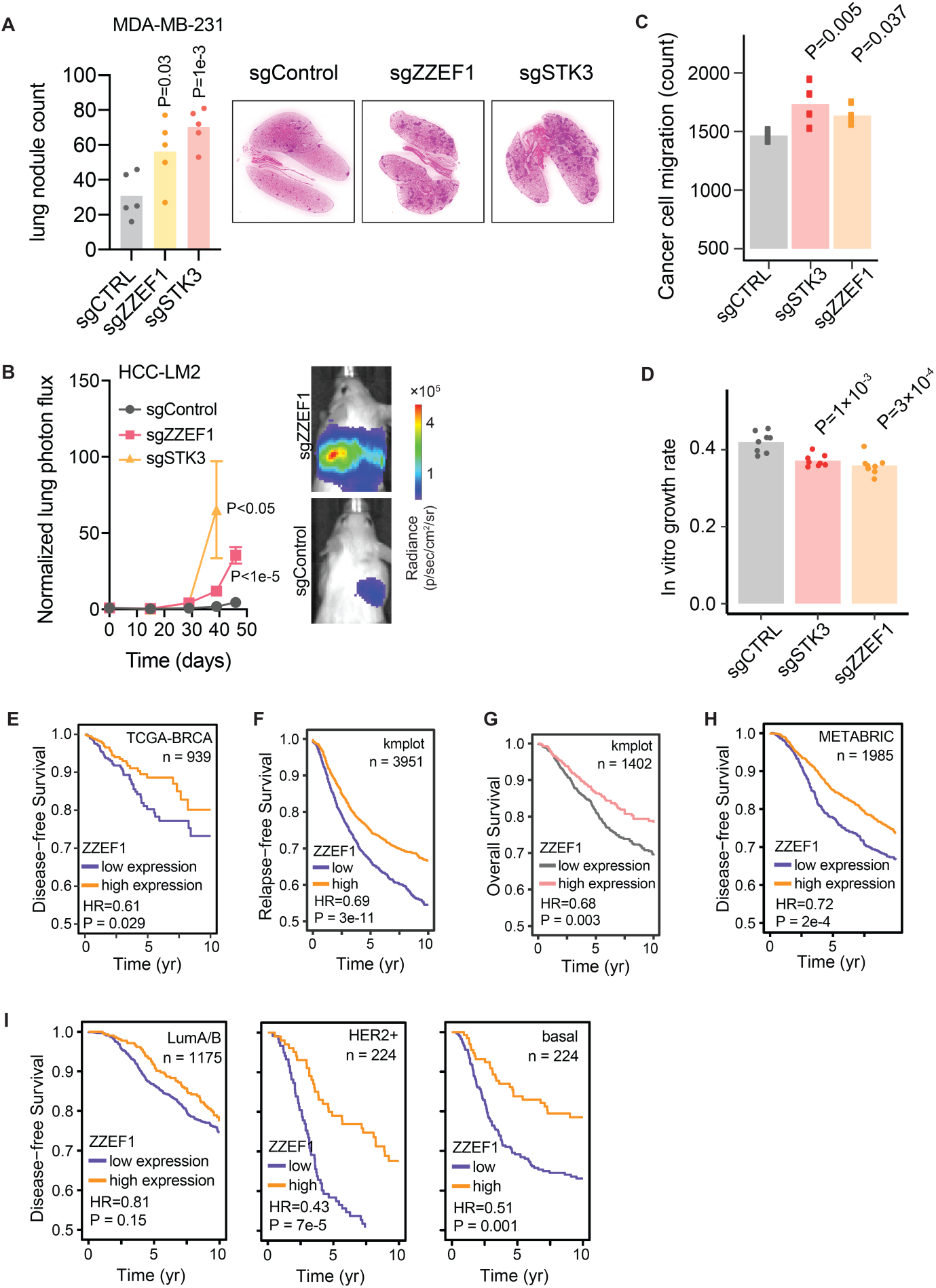
ZZEF1-tRNA-Lys^UUU^-STK3 deficiency drives breast cancer metastasis and migration. **(A)** Gross histology of extracted lungs showing significantly increased metastatic burdens after ZZEF1 and STK3 knockdowns compared to control cells. *P*-values were calculated using *t*-tests. **(B)** Bioluminescence imaging showing lung colonization in HCC1806-LM2 cells with ZZEF1 knockdown and STK3 knockdown compared to control cells. **(C)** Migration assay showing enhanced migration capability of MDA cells with ZZEF1 knockdown and STK3 knockdown compared to control cells. *P*-values were calculated using *t*-tests. **(D)** Proliferation assay showing reduced growth rates of MDA cells with ZZEF1 knockdown and STK3 knockdown compared to control cells. *P*-values were calculated using *t*-tests. **(E)** Kaplan-Meier survival curve illustrating the correlation between tumor ZZEF1 levels and disease-free survival in a collection of breast cancer patient cohorts in TCGA dataset. **(F)** Similar to **(E)**, but showing the correlation between ZZEF1 levels and relapse-free survival. **(G)** Similar to **(E)**, but showing the correlation between ZZEF1 levels and overall survival. **(H)** Similar to **(E)**, but showing the correlation with other public datasets. **(I)** Kaplan-Meier survival curves illustrating the correlation between tumor ZZEF1 levels and disease-free survival in breast cancer subtypes, including LumA/B, HER2+, and basal subtypes.

**Figure S5.**
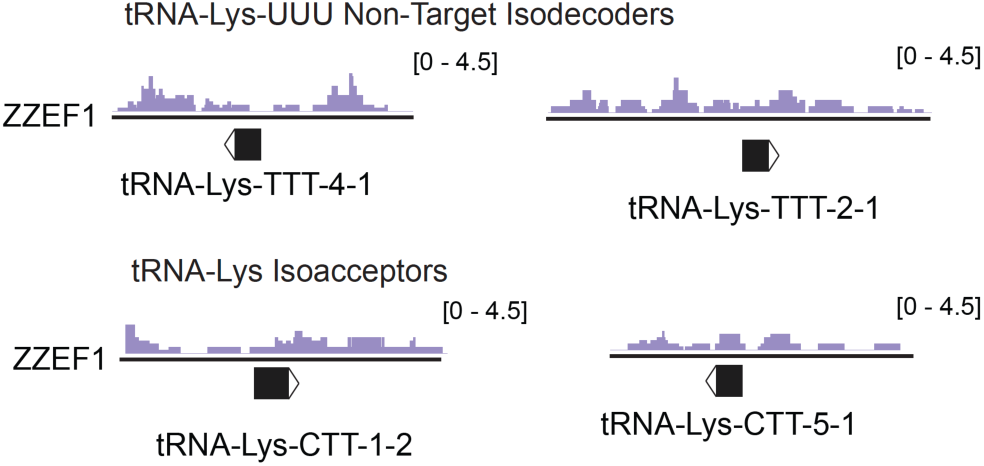
**ZZEF1 binding profiles in non-target loci.**

**Figure S6.**
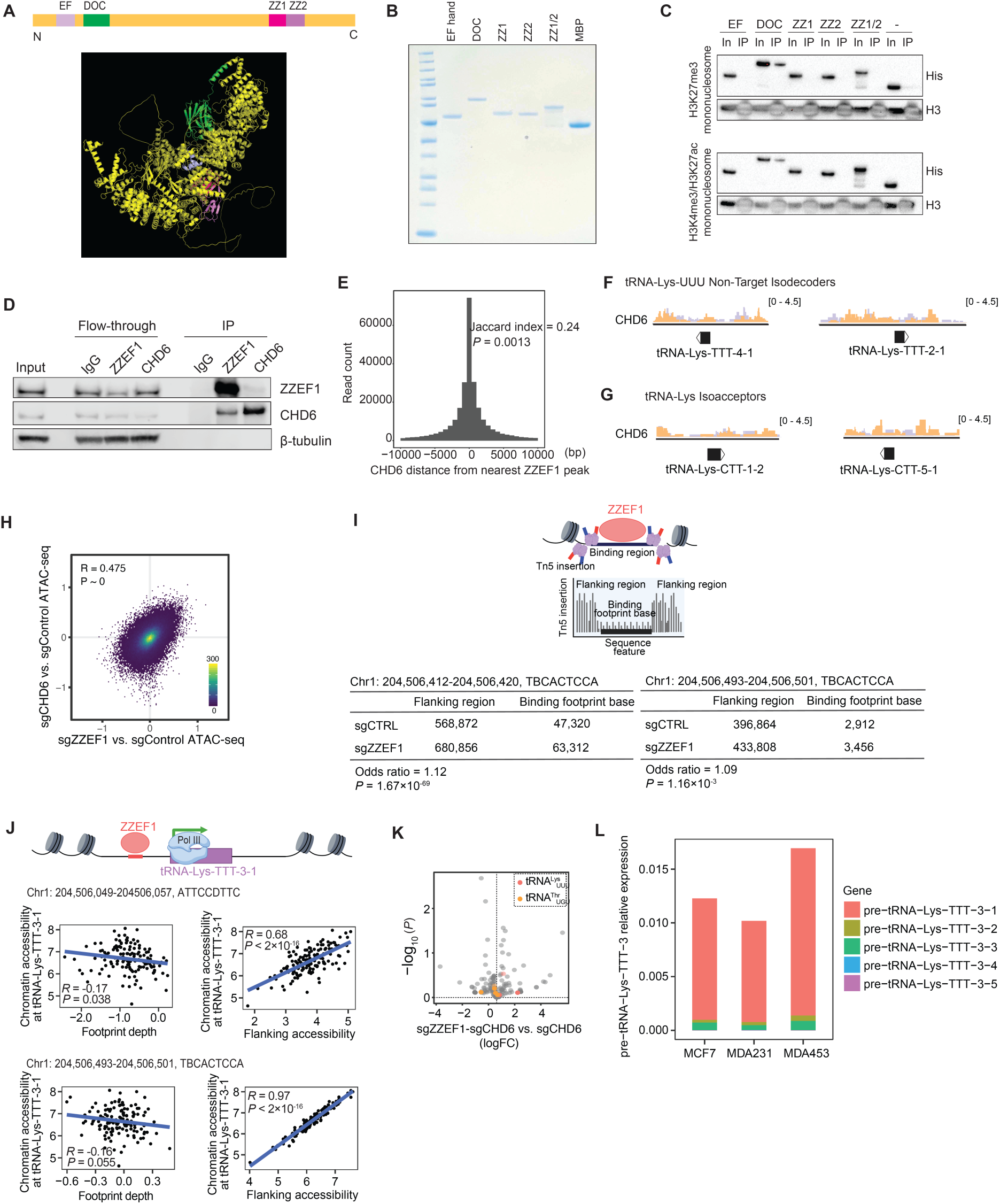
ZZEF1 and CHD6 work together in proximal binding and function in chromatin remodeling. **(A)** Schematic representation of ZZEF1 domains. EF hand (111-146 amino acid), DOC (226-405 amino acid), ZZ1 (1778-1833 amino acid), ZZ2 (1827-1882 amino acid), ZZ1/2 (1778-1882 amino acid). **(B)** PAGE analysis of purified ZZEF1 domains, including EF hand, DOC, ZZ1, ZZ2, the combined ZZ1/2, and MBP (control). **(C)** Pulldown immunoprecipitation (IP) results showing interactions between ZZEF1 domains and mononucleosomes with H3K27me3 or H3K4me3/H3K27ac modifications. In, input. IP, biotin pulldown. **(D)** Co-immunoprecipitation (co-IP) and Western blot analysis showing protein abundance of *β*-tubulin, CHD6, and ZZEF1 in input, IgG control, ZZEF1 IP, CHD6 IP, and flow-through fractions. **(E)** Histogram showing a high degree of overlap between CHD6 and ZZEF1 binding peaks on a genome-wide scale. *P* value was calculated with a permutation test. **(F)** Normalized ChIP-seq tracks showing enriched CHD6 binding (orange) in the proximal regions of non-target loci using a CHD6-specific antibody, compared to an IgG control in MDA cells. The purple tracks represent ZZEF1 binding from Fig. 5B. **(G)** Similar to **(F)**, but showing results at tRNA-Lys isoacceptor loci. **(H)** Density plot showing a positive correlation between chromatin accessibility changes after CHD6 knockdown and chromatin accessibility changes after ZZEF1 knockdown compared to control cells. **(I)** Two by two tables illustrating the significant reduction in ZZEF1 binding potential at the tRNA-Lys-TTT-3 loci following ZZEF1 knockdown. *P* values were calculated with fisher exact tests. **(J)** Scatter plot showing the association between reduced binding potential at ZZEF1-specific sequence contexts and decreased chromatin accessibility at tRNA-Lys-TTT-3-1 locus across breast cancer samples. High footprint depth indicates low binding potential. Positive correlation is also observed between chromatin accessibility at tRNA-Lys-TTT-3-1 locus and flanking regions of ZZEF1-specific sequence contexts in TCGA breast cancer patient data. Pearson *R* and associated *P*-value are shown. **(K)** Volcano plot showing alterations in mature tRNA expression profiles between CHD6 knockdown MDA cells and CHD6 knockdown cells in a ZZEF1-depleted background. **(L)** Bar plot showing pre-tRNA-Lys-TTT-1/2/3/4/5 relative expression in MCF7 cells, MDA231 cells and MDA453 cells.

